# Structure determinants of DANGEROUS MIX 3, an alpha/beta hydrolase, for triggering NLR-mediated genetic incompatibility in plants

**DOI:** 10.1101/2024.10.28.620575

**Authors:** Gijeong Kim, Wei-Lin Wan, Nayun Kim, Yi Yun Tan, Nuri Charoennit, Rachelle R. Q. Lee, Yin Yin Liew, Shen Kai Ng, Yizhong Zhang, Ji-Joon Song, Eunyoung Chae

**Affiliations:** Department of Biological Sciences, KI of BioCentury, Korea Advanced Institute of Science and Technology (KAIST), Daejeon, 34141, Korea; Department of Biological Sciences, National University of Singapore, 117558, Singapore; Department of Plant and Microbial Biology (IPMB), Zurich-Basel Plant Science Center, University of Zurich, Zurich, 8008, Switzerland (current); Department of Otolaryngology, Yong Loo Lin Scholl of Medicine, National University of Singapore, 117545, Singapore (current); Department of Medicine, Yong Loo Lin Scholl of Medicine, National University of Singapore, 119228, Singapore (current); Department of Biology, University of Oxford, Oxford, OX1, UK (from Jan 2025)

**Keywords:** Genetic incompatibility, NLR, autoimmunity, alpha/beta hydrolase, cryo-EM, structure, oligomer, hexamer

## Abstract

Genetic incompatibility occurs when a mismatched pair of plant immune components mounts autoimmune responses in hybrids. Highly diversified NLR receptors are main culprits of the genetic conflict, recognizing host proteins from different origin as immune trigger. Here, we report the molecular mechanism underlying a DANGEROUS MIX (DM) autoimmunity, comprising DM2h/RPP1 NLR and its incompatible partner DM3, an alpha/beta hydrolase. Cryo-electron microscopy reveals the oligomeric nature of two natural DM3 variants in a trimer of dimer configuration. The polymorphism triggering autoimmunity is located at the dimer interface, resulting in drastic structural differences such that dimerizing helix and loop reinforcing the interface is lost and disordered. Structure-function analysis shows that integrity of the dimer interface, but neither maintenance of hexamer nor its enzymatic activity, is the key factor contributing to autoimmunity. Our finding pinpoints checkpoints embedded in the oligomeric configuration of a host enzyme that controls the switching mechanism of NLR activity.

## INTRODUCTION

Hybrid necrosis is a phenomenon often observed in the first and later generation of hybrid plants suffering from unnecessary mounting of immune responses and accompanying developmental defects. This form of genetic incompatibility has been actively studied in the past decade, revealing numerous incompatible genetic elements enriched with immune genes. Traits associated with the causal incompatible genes include extensive polymorphism, genomic instability and elevated mutation rate, which have presumably contributed to diversifying immune receptor repertoires in natural populations. However, evolutionary consequences of receptor diversification come at the cost of immune system incompatibilities. Previous systematic species-wide studies of hybrid necrosis in *Arabidopsis thaliana* (herein Arabidopsis) uncovered that multiple system failure points are largely attributed to mismatched immune components involving members of the nucleotide-binding domain and leucine-rich repeat (NLR) intracellular immune receptors^1–5^. In hybrid necrosis plants, the process of mounting immune responses by an NLR is misregulated^3,6^, as the receptor evolved from one parent recognizes a host molecule from the other parent as a trigger. Thus, pairs of incompatible immune genes offer an opportunity to study NLR activation mechanisms as well as biological processes being monitored by NLRs.

With a modular tripartite domain structure, NLRs have evolved to monitor cellular homeostasis under pathogenic attacks^7^. Only a small fraction of functionally annotated plant NLRs has been assigned to matching pathogenic effectors that are secreted into plant cells upon invasion^8,9^; most NLRs in plants are thought to perceive molecular disturbance of certain host proteins, which are formally termed as guardees or decoys^10–13^. NLR activation by this indirect recognition of modified-self has been demonstrated with several NLRs, including Arabidopsis RPM1, RPS2 and RPS5, which recognize post-translational modification status of host proteins under their surveillance as the activation cues^14–20^. The mechanistic understanding of NLR activation in plants is facilitated by recent structural determination of oligomeric NLRs with cryo-electron microscopy (cryo-EM), including RPP1 and Roq1 as tetrameric complexes bound to their respective effectors, and ZAR1 as a pentameric complex bound to decoy molecules activated by an effector^21–24^. Full NLR activation results in acute immune responses in plants, leading to substantial transcriptional reprogramming to confer disease resistance and often immunological cell death termed as the hypersensitive response (HR) ^25^. Recent studies show that a particular subfamily of plant NLRs carrying the Toll/interleukin-1 receptor (TIR) domain at the N-terminus engages small molecules for concomitant immune signaling^26–32^. Oligomerization of TIR-containing NLRs enables the assembly of catalytic core surrounding the conserved glutamic acid residue, which is critical for the production of various types of purine nucleotides through enzymatic activities of the TIR domains. Subsequent binding of such small molecules to core immune signaling complexes, including EDS1 and their partners belonging to the alpha/beta hydrolase (ABH) superfamily, then activates helper NLRs for executing immune responses.

Although the process of mounting immune responses by an NLR should be well regulated for successful defence, studies of autoimmunity showed that numerous plant NLRs are prone to activate upon slight changes in the genetic background either introduced by transgenes, mutations or hybridization^33–36^, hinting at single-point failures in the plant immune system lying at the receptor level. This also indicates that there is a wide range of molecular processes that NLRs monitor, only a fraction of which could have been assigned with guardee/decoy functions due to the limited number of lab-amenable pathosystems. Several episodes of inadvertent NLR activation are genetically and molecularly well characterized as hybrid necrosis. Such NLRs and their incompatible interacting partners are designated as DANGEROUS MIX (DM) in Arabidopsis^3,4^. The hypervariable *DM2* locus, encoding RPP1-like NLRs as a multigene cluster, stands out as a recurrent source of autoimmune-risk alleles in natural populations^4^. Additionally, *DM2h* from the widely used accession L*er* is known to trigger autoimmunity by three independent causes, including a natural polymorphism in STRUBELLIC RECEPTOR FAMILY 3 (SRF3), a receptor kinase that has recently been reported to sense environmental iron levels^37,38^, a mutation in O-acetylserine(thiol)lyase (OASTL), a cysteine synthase^39,40^, and mislocalization of EDS1^5,41^. Given the versatile ways of making DM2/RPP1 NLR uncontrolled by homeostasis disturbance in the cell, the underlying molecular mechanism for DM2 activation by host factors, together with structural knowledge gained from effector-driven RPP1 activation^21^, is likely to provide us with comprehensive understanding on NLR activation.

Here, we focus on the genetic interaction between *DM2h,* encoding an RPP1-like NLR, and *DM3,* encoding a prolyl aminopeptidase belonging to the ABH superfamily, to study NLR activation by a host enzyme. A specific allele of *DM3, DM3*^Hh-0^, triggers autoimmunity in the presence of *DM2h*^Bla-1^, an NLR allele^4^. We aimed to investigate how this particular variant of host enzyme DM3 activates immunity with DM2h^Bla-1^ by determining the cryo-EM structures of DM3^Hh-0^ and DM3^Col-0^, which is a variant that does not cause autoimmunity with DM2h^Bla-1^. Our structural determination and biochemical characterization unveil multiple interaction interfaces of DM3, conditioning various oligomeric statuses *in vitro* and *in vivo*, and a particular polymorphism-mediated structural disruption of the DM3 oligomers that leads to activate NLR-mediated immunity. In addition, we were able to discern the immune-activating function of DM3 from its catalytic activity. Our work illustrates an evolutionary strategy of the multigene *DM2/RPP1* NLR cluster to generate an allele to recognize structural integrity of the host enzyme in addition to conventional effector sensors, expanding its guarding units to cover cellular homeostasis in check.

## RESULTS

### Natural polymorphism of DM3 responsible for immune activation by DM2h NLR

Previously, DM3 was annotated as a prolyl aminopeptidase (PAP2 encoded by At3g61540), capable of catalytically cleaving a proline residue located at the N-terminal end of a peptide^42^. DM3 carries a typical catalytic triad consisting of S214, H490, and D462 with a predicted transit peptide (TP) of 52 amino acids at the N-terminus (Figure 1A). Phylogenetic analysis demonstrated that DM3 forms a distinct clade with a group of homologs from other plant species, while the other functionally annotated PAP from Arabidopsis (PAP1: At2g14260) ^43^ is found in a separate clade (Figure 1B). Protein sequence alignment showed that the DM3 homologs are characterized with an additional domain inserted within the ABH fold (Figures 1A and S1). Before the annotation of *PAP2*, we identified *DM3* as a partner of the NLR gene *DM2/RPP1*^4^. As the causal variant of DM3 from Hh-0 for hybrid necrosis harbours two amino acid differences (T165I and T359A) from that of Col-0 (Figure 1A), we surveyed the polymorphism spectrum of DM3 from the Arabidopsis 1001 genome database^44^. From 1135 accessions, we identified 37 representative DM3 protein sequences and generated a haplotype network. The majority of accessions encode DM3 proteins identical to that in Col-0, with 27 haplotypes differing from DM3^Col-0^ by no more than two amino acids (Figure 1C). The hybrid necrosis variant DM3^Hh-0^ differs at two positions from DM3^Col-0^; the rare variation T165I is found only in Hh-0 on top of the polymorphism T359A that is found in 21 accessions (Figure 1C). The allele-specific effect triggering DM2h^Bla-1^-dependent immune responses can be recapitulated as HR in the heterologous *Nicotiana benthamiana* (*Nb*) system^4^ (Figure 1D). Previously, Ghifari and colleagues reported that the N-terminal TP is functional in mitochondrial targeting, which we noted to interfere with further biochemical studies in terms of protein expression. We sought to test whether the immune activation of DM2h^Bla-1^ with DM3^Hh-0^ requires the first 52 amino acids of DM3. Co-infiltration of HA-_ΔTP_DM3^Hh-0^ with DM2h^Bla-1^ in *Nb* resulted in robust and consistent HR, which was comparable to that of full-length DM3^Hh-0^. Meanwhile, TP deletion did not render DM3^Col-0^ the capacity of triggering HR with DM2h^Bla-1^ (Figure 1D). The finding of TP not contributing to the immune function of DM3 suggests that cytoplasmic localization of DM3 proteins is sufficient to activate the intracellular DM2h NLR. Subsequent western blotting demonstrated that even the constructs encoding the full-length DM3 variants produced the major detectable forms in the SDS-PAGE equivalent to those of the TP-truncated DM3 proteins (Figure 1E). Our data indicates that _ΔTP_DM3 proteins retain original immune functions and thus qualify as functional units suitable for further biochemical investigation.

**Figure 1.**
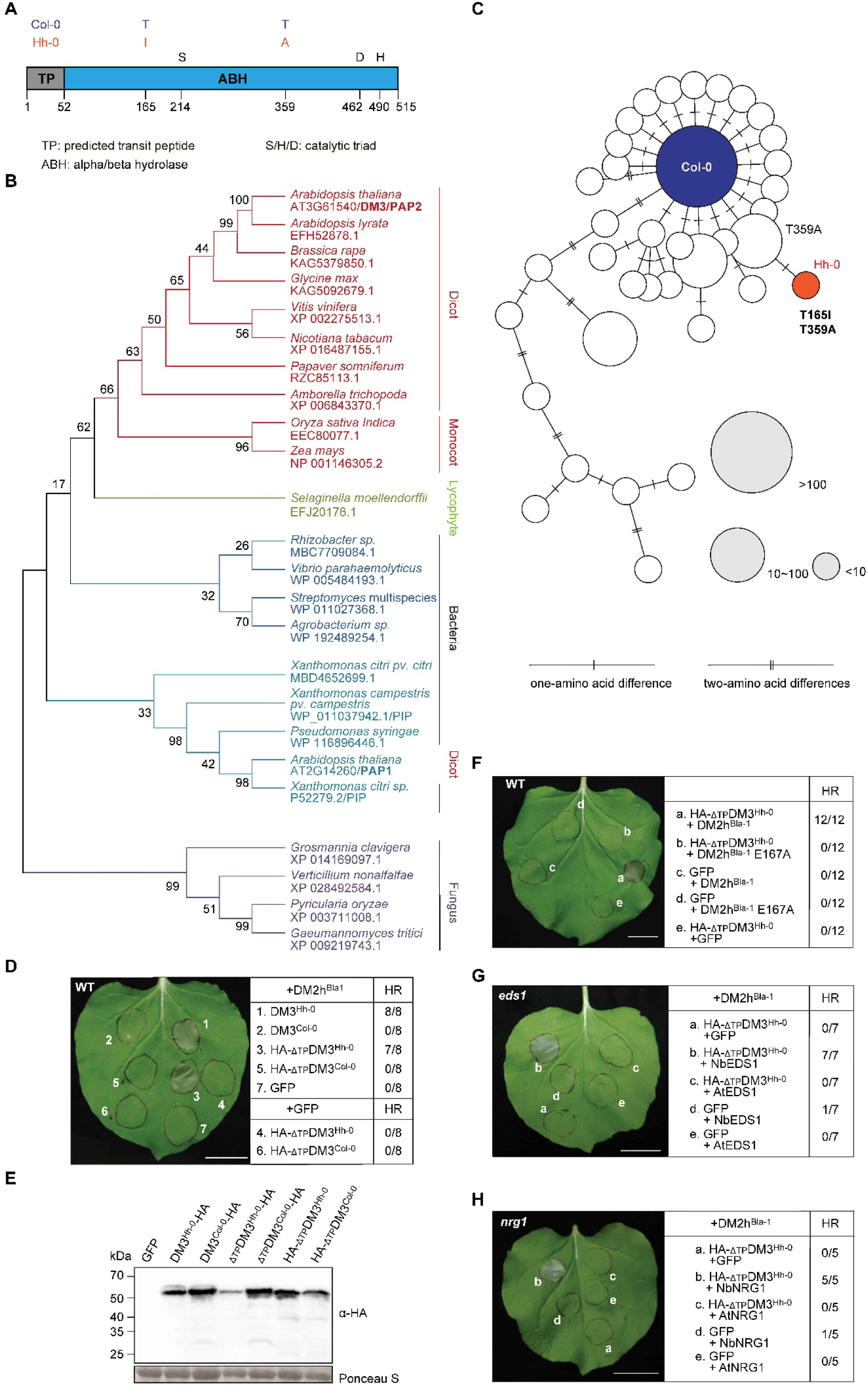
Natural variation of DM3 and its allele-specific immune function with DM2h NLR. (A) Schematic diagram of DM3 protein. Two amino acid differences between DM3^Col-0^ and DM3^Hh-0^, a predicted transit peptide (light grey), an alpha/beta hydrolase domain (blue) and a catalytic triad are shown. (B) A maximum-likelihood tree of DM3 homologs. The values besides the branches represent percentage bootstrap support. The species and GenBank or NCBI reference numbers used for phylogenetic analysis are indicated. (C) DM3 protein haplotype network of 37 distinct DM3 alleles in Arabidopsis generated by R package, pegas. The three different sizes of circles indicate the number of accessions with a given haplotype. (D) Expression of DM3 variants with DM2h^Bla-1^ or GFP in *Nb*. DM2h^Bla-1^ and full-length DM3 were driven by endogenous promoters, while DM3 without predicted transit peptide and GFP were driven by 35S promoters. Photos were taken five days after infiltration. Numbers indicate the incidents of confluent HR out of an experiment with eight leaves. Scale bar equals 2 cm. (E) Western blot of DM3 variants transiently expressed in *Nb*. ΔTP indicates the deletion of a transit peptide spanning the first 52 amino acids. Expression was driven by the 35S promoter, and leaf samples were collected at 28 hours after infiltration. (F) Expression of DM3^Hh-0^ with DM2h^Bla-1^ or DM2h^Bla-1^ E167A in *Nb*. DM2h^Bla-1^ and DM2h^Bla-1^E167A were driven by endogenous promoter; DM3^Hh-0^ and GFP by 35S promoter. (G) Expression of DM3^Hh-0^, DM2h^Bla-1^ with EDS1 in *Nb-eds1* mutant line. NbEDS1 was driven by endogenous promoter; AtEDS1 was driven by 35S promoter. (H) Expression of DM3^Hh-0^, DM2h^Bla-1^ with NRG1 in *Nb nrg1* mutant line. NbNRG1 was driven by endogenous promoter; AtNRG1 was driven by 35S promoter. Predicted size of DM3-HA: 64 kDa; HA-_ΔTP_DM3/_ΔTP_DM3-HA: 58.3 kDa. See also Figure S1.

Further characterization of DM2-_ΔTP_DM3 autoimmunity was carried out using the *Nb* system to define downstream signaling pathways. When the conserved, catalytic glutamic acid residue was mutated in the TIR-domain of DM2h NLR (E167A), HR was completely abrogated (Figure 1F). HR was not observed when the paired *DM* constructs were infiltrated in the *Nb* mutants of *eds1* and of *nrg1*, a helper NLR, while the HRs reappeared when the infiltration was supplemented with *EDS1* or *NRG1* cloned from *Nb* but not from Arabidopsis (Figures 1G and 1H). These HR assays indicate that the autoimmunity conforms to the canonical signaling pathways that most known TIR-containing NLRs engage in to activate effector-triggered immunity.

### The cryo-EM structure of DM3^Col-0^ as a trimer of dimers

To dissect the molecular basis of the NLR-mediated immune activation by DM3, we first expressed N-terminally truncated DM3 (_ΔTP_DM3; 52-515) to produce recombinant DM3^Col-0^ proteins from *E. coli.* The recombinantly purified _ΔTP_DM3^Col-0^ protein was eluted at high molecular weight size fraction (158∼440 kDa) in a gel-filtration chromatography, suggesting that _ΔTP_DM3^Col-0^ (51.9 kDa as a monomer) forms an oligomer (Figure 2A). Further examination of the DM3 oligomer by negative stain EM indicated that _ΔTP_DM3^Col-0^ forms a homogeneous large oligomer complex approximately 100 Å in size (Figure 2B). To determine the molecular architecture of the DM3 oligomer, _ΔTP_DM3^Col-0^ protein structure was examined by cryo-EM. From the collected 4,055 cryo-EM micrographs, 330,509 final particles were used for reconstructing a cryo-EM map of _ΔTP_DM3^Col-0^ at 3.39 Å resolution (Figures 2C and S2). With no symmetry imposed during the image processing, the cryo-EM map of _ΔTP_DM3^Col-0^ shows a hexameric complex 108 Å in diameter and 70 Å in depth (Figure 2C). The cryo-EM map clearly resolved the side chain density, and we built an atomic model of _ΔTP_DM3^Col-0^ (68-515) *de novo* (Figure S2 and Table S1). The cryo-EM structure of _ΔTP_DM3^Col-0^ revealed a hexameric complex composed of a trimer of _ΔTP_DM3^Col-0^ dimers (Figure 2D). The six _ΔTP_DM3^Col-0^ subunits are arranged in a D3 symmetry with three-fold rotational axis that penetrates a centre hole of the hexamer and two-fold rotational axes at the side of the hexamer (Figure 2D).

**Figure 2.**
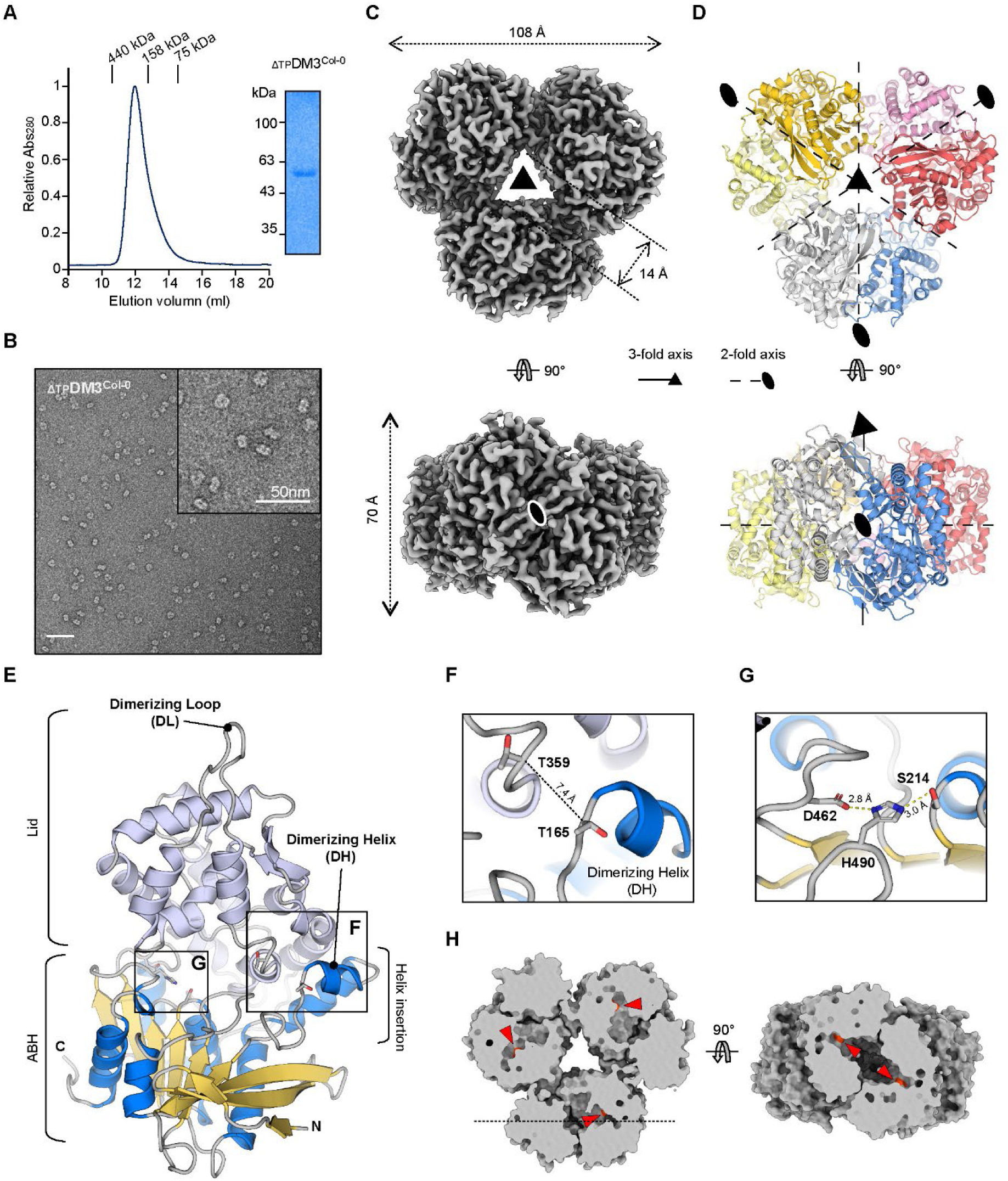
Biochemical characterization and cryo-EM structure of _ΔTP_DM3^Col-0^. (A) A size-exclusion chromatography on the recombinant _ΔTP_DM3^Col-0^ protein shows its oligomeric status *in vitro*. SDS-PAGE shows a purity of the _ΔTP_DM3^Col-0^ protein at the high molecular weight fraction. (B) A negative stain EM shows the shape of the _ΔTP_DM3^Col-0^ oligomer. An enlarged micrograph is shown in the right corner. The scale bar represents 50 nm (C) A 3.39 Å resolution cryo-EM map of _ΔTP_DM3^Col-0^ is presented in grey. (D) The cryo-EM structure of _ΔTP_DM3^Col-0^ hexamer is presented as a cartoon. Six _ΔTP_DM3^Col-0^ subunits are differentially colored. A three-fold rotational axis (triangle) and two-fold (oval) rotational axis are labelled on the cryo-EM map and structure of _ΔTP_DM3^Col-0^ hexamer. (E) A cryo-EM structure of _ΔTP_DM3^Col-0^ monomer is shown in a ventral view (from the core cavity of the hexamer). ɑ-helices and β-sheets of ABH domain are colored blue and yellow. The secondary structures of the lid domain are colored pale purple. N- and C-terminus of _ΔTP_DM3^Col-0^ are labelled, respectively. The Dimerizing Helix (DH) and Dimerizing Loop (DL) are indicated. (F) T165 and T359 of DM3^Col-0^ are located in close proximity at three-dimensional space. The distance between Cɑ atoms of T165 and T359 is indicated with a dotted line. (G) A detailed view of the catalytic triad is illustrated. Electrostatic interactions within the catalytic triad are shown in dotted lines. (H) A slice of top (left) and side (right) view of _ΔTP_DM3^Col-0^ hexamer visualizes the catalytic active sites. S214 is colored red and pointed by red arrowheads. The horizontal dashed line indicates a slab to visualize the catalytic triad with red arrowheads. See also Figure S2, S3 and Table S1.

To further elucidate the molecular details of the hexamer structure, we analysed the monomer structure of DM3^Col-0^. Having an identical conformation, each DM3^Col-0^ monomer has a typical ABH domain structure, in which a large lid domain is inserted spanning amino acid 244-442^45^ (Figures 2E and S3). The ABH domain has a canonical fold that has eight β strands flanked by several α helices^46^, and the lid domain interrupts the ABH domain between β6 strand and α4 helix (Figures S3A and S3B). In addition, a helix insertion (154 - 186) in the ABH domain interacts with the lid domain, structurally forming a single domain. (Figures 2E and S3A). Interestingly, the cryo-EM structure shows that the polymorphic residues of DM3^Col-0^, T165 and T359, are clustered at the interface between the helix insertion and the lid domain (Figures 2F and S3A). The catalytic triad (S214-H490-D462) is located at a distance from T165 and T359, and the hexamer structure compartmentalizes the catalytic triads within the hexameric chamber (Figures 2G and 2H). The cryo-EM structure of DM3^Col-0^ delineates the oligomeric state of DM3 and suggests a functional separation of the HR triggering activity of DM3^Hh-0^ from the SHD-dependent catalytic activity of DM3.

### Dimer and trimer interfaces within the DM3 hexameric structure

As the DM3^Col-0^ hexamer is configured as a trimer of dimeric units (Figure 2D), we analysed the inter-molecular interactions contributing to the hexamer formation of DM3. At an interaction interface within the dimer (herein dimer interface), a long loop connecting α4 and α5 helices in the lid domain interacts with the α2-1 and α2-2 helix insertion of the other subunit (Figures 3A, S3A, and S3B). We named the loop and helix as Dimerizing Loop (DL; 329-344) and Dimerizing Helix (DH; 166-171), respectively (Figures 2E, 3B, S3A, and S3B). At the dimer interface, the extended DL is stabilized by the interaction with the DH. Specifically, Q171 forms hydrogen bonds with the main chain of the DL (Figure 3B). L335 and V336 of the DL form hydrophobic interactions with L164, F172, and Y184 of the helix insertion (Figure 3B). Interestingly, one of the polymorphic residues, T165, is located at the beginning of the DH and participates in an electrostatic interaction with S167 and D332 (Figure 3B and 3C). The other polymorphic residue, T359, is located in proximity to T165 and forms an electrostatic interaction with D358, which in turn interacts with N349 of the lid domain in the other DM3 molecule (Figure 3C). A head-to-tail interaction at the dimer interface leads to a 2-fold symmetry, which in turn forms an inter-dimer interface (herein trimer interface) to complete the hexamer complex (Figure 3A). The trimerization of DM3^Col-0^ dimers is mediated by the interaction between the C-terminus and ABH domain of DM3 (Figure 3D). A close look into the trimer interface revealed that five C-terminal residues (KKPLF; 511-515) are buried in a pocket between the lid and ABH domain of the adjacent DM3^Col-0^ dimer (Figure 3E). This junction renders the dimers to form a salt bridge between the C-terminal carboxyl group of F515 and the ε-amino group of K266 as well as hydrophobic interactions between F515 and M489, L514 and the aliphatic side-chain of K266, and P513 and P79 (Figure 3E). At the trimer interface, two ABH domains form an antiparallel β sheet using their β8 strands. Above the β sheet, Y460, F467, and V484 from each ABH domain form hydrophobic interactions (Figure 3F). Detailed structural analysis of molecular interactions in _ΔTP_DM3^Col-0^ hexamer revealed the significance of the polymorphic residues at the dimer interface.

**Figure 3.**
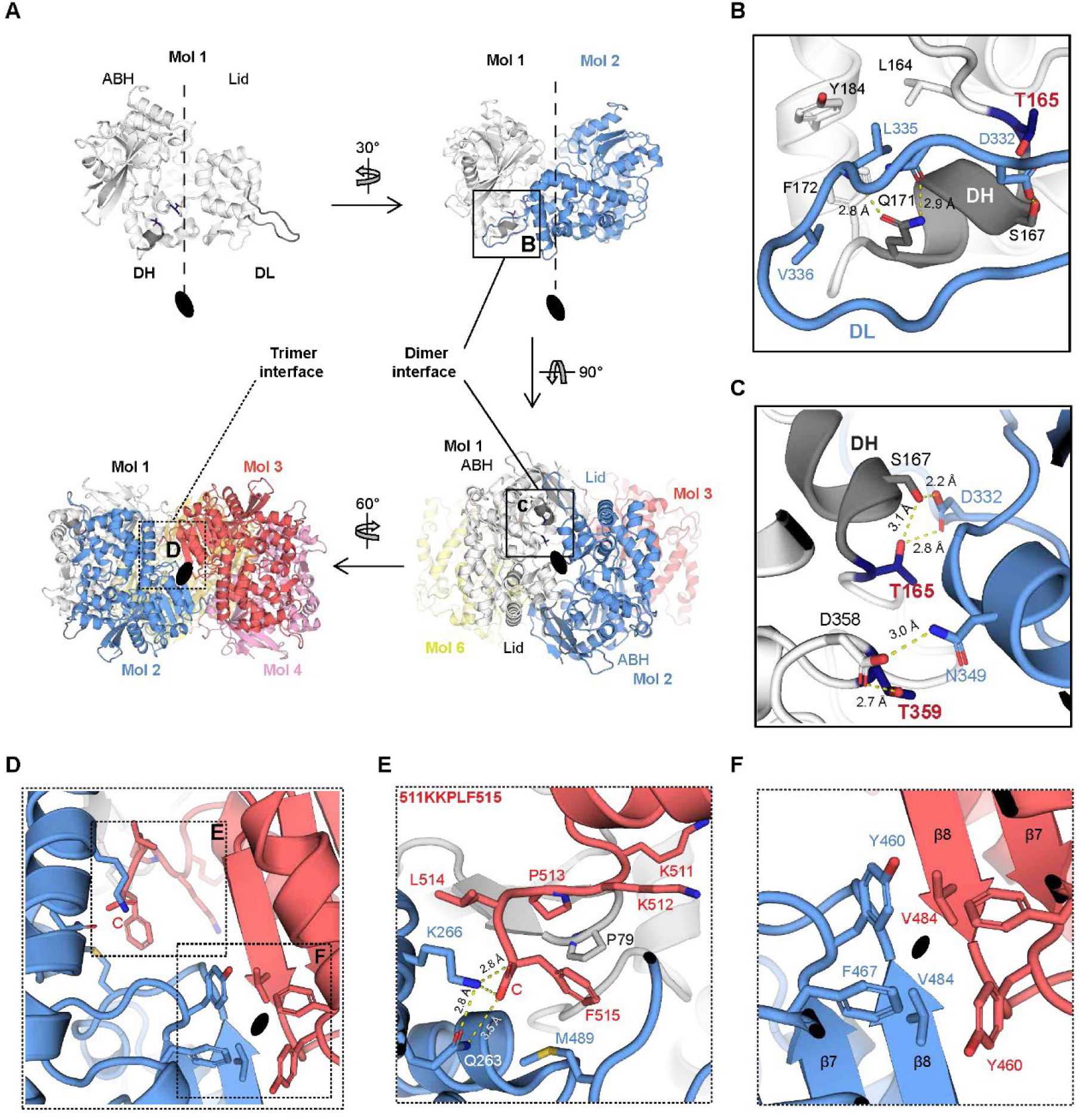
Characteristics of the dimer and trimer interfaces of DM3^Col-0^ hexamer. (A) The dimer and trimer interface of DM3^Col-0^ are specified on the cryo-EM structure of _ΔTP_DM3^Col-0^ monomer, dimer and hexamer. The dimer interface formed by Mol 1 and Mol 2 is outlined by a box and the trimer interface formed by two _ΔTP_DM3^Col-0^ dimers is indicated in a dotted box. (B) The dimer interface formed by DH and DL is shown in detail. DH and T165 from Mol 1 are colored with dark grey and navy, respectively. (C) The dimer interface near to T165 and T359 (navy) is shown in detail. T165, T359 and some interacting residues in the dimer interface. (D) The trimer interface shows C-terminus in dotted box (E) and the hydrophobic residues located at the two-fold rotational axis in dotted box (F). (E) A detailed view of the C-terminus of _ΔTP_DM3^Col-0^ is depicted. The carboxyl group of F515 is denoted by C. (F) The anti-parallel β sheet formed by ABH domains from two _ΔTP_DM3^Col-0^ dimers is shown with the hydrophobic amino acids. Residues of Mol 1, Mol 2, and Mol 3 are labelled in black, blue and red. The electrostatic interactions are depicted by dotted lines. The horizontal dashed line indicates a slab to visualize the catalytic triad with red arrowheads. The sliced view is shown in right side. See also Figure S3.

### T165I polymorphism-mediated disruption of the dimer interface in DM3^Hh-0^

To dissect the molecular consequences of the two-residue polymorphism found in DM3^Hh-0^ on structural and biochemical properties, we expressed and purified recombinant _ΔTP_DM3^Hh-0^ from *E. coli*. Like its Col-0 counterpart, the purified _ΔTP_DM3^Hh-0^ recombinant protein was eluted at the high molecular weight fraction from gel-filtration and the large oligomer was observed by negative stain-EM like _ΔTP_DM3^Col-0^, suggesting that the polymorphism does not affect the integrity of the DM3 hexameric structure *in vitro* (Figures S4A-S4C). To elucidate the molecular architecture of DM3^Hh-0^, we determined a cryo-EM structure of _ΔTP_DM3^Hh-0^, following similar procedures used for _ΔTP_DM3^Col-0^. A total of 14,531 micrographs of _ΔTP_DM3^Hh-0^ were collected and 413,056 particle images were used for a cryo-EM map reconstruction of _ΔTP_DM3^Hh-0^ at 2.89 Å resolution (Figure S4D). With clearly resolved side chain density, we built an atomic model of _ΔTP_DM3^Hh-0^ in the cryo-EM map (68-515) (Figure 4A, S4, and Table S1). The global structure of DM3 hexamer is conserved between _ΔTP_DM3^Hh-0^ and _ΔTP_DM3^Col-0^ with RMSD value of 0.796 Å (Figure 4B). However, at the dimer interface, the polymorphisms of _ΔTP_DM3^Hh-0^ (I165 and A359) lead to a drastic structural change in DL and DH in contrast to the ordered interaction between DH and DL of _ΔTP_DM3^Col-0^ (Figures 4C and 4D).

**Figure 4.**
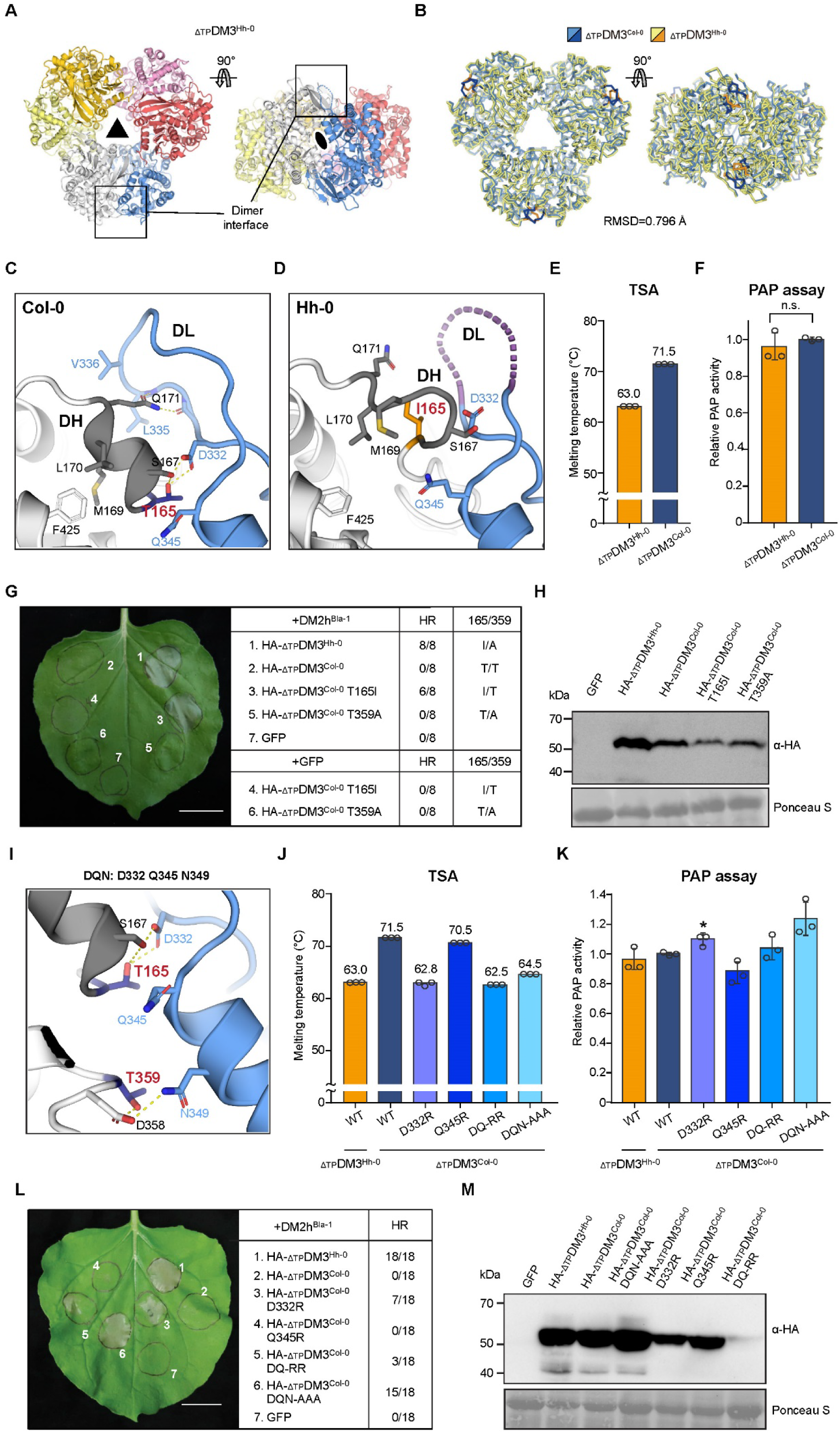
Cryo-EM structure of _ΔTP_DM3^Hh-0^ and structure-guided dimer interface mutants of DM3^Col-0^ show polymorphism-mediated structural change and immune activation. (A) A cryo-EM structure of _ΔTP_DM3^Hh-0^ hexamer is displayed. Each chain is colored with different colors. The dimer interface of _ΔTP_DM3^Hh-0^ is outlined in a box. (B) The cryo-EM structures of _ΔTP_DM3^Col-0^ and _ΔTP_DM3^Hh-0^ are depicted by ribbon and superimposed with each other with RMSD value of 0.796 Å. _ΔTP_DM3^Col-0^ is colored blue and _ΔTP_DM3^Hh-0^ yellow. The DH and DL of _ΔTP_DM3^Col-0^ and _ΔTP_DM3^Hh-0^ are colored navy and orange respectively. Type or paste caption here. (C) The dimer interface of _ΔTP_DM3^Col-0^ near T165 (navy). DH is colored with dark grey. (D) The dimer interface of _ΔTP_DM3^Hh-0^ is shown near I165 (orange) with the same view of (C). The DH is melted and the disordered region of DL (335-340 a.a.) is colored in purple. (E) Thermal shift assay (TSA) shows the melting temperature of _ΔTP_DM3^Hh-0^ (orange) and _ΔTP_DM3^Col-0^ (navy). Each data point is represented by a circle (n = 3). (F) The enzymatic activity assay shows normalized prolyl aminopeptidase (PAP) activity of _ΔTP_DM3^Col-0^ and _ΔTP_DM3^Hh-0^. Individual data points (n = 3) and standard deviation are represented. n.s. represents no significant difference. (G) Expression of DM3 variants with DM2h^Bla-1^ or GFP in *Nb*. The expression of DM2h^Bla-1^ was driven by endogenous promoters, while _ΔTP_DM3s and GFP were driven by 35S promoter. Photos were taken five days after infiltration. HR success rate is indicated with the number of leaves that shows fully developed HR. The scale bar equals 2 cm. (H) Western blot of DM3 variants. *Nb* leaf samples were collected at 28 hours after infiltration. (I) The dimer interface of DM3^Col-0^ shows polar amino acids (D332, Q345 and N349; herein D, Q and N) on the opposite side of T165 and T359. (J) TSA shows melting temperatures of purified dimer interface mutant proteins (DQ-RR; D332R Q345R and DQN-AAA; D332A Q345A N349A). Each data point is represented by a circle (n = 3). (K) The enzymatic activity assay shows normalized PAP activity of the dimer interface mutants. Individual data points (n = 3) and standard deviation are represented, and asterisk indicates p-value from two-tailed t-test between wild type _ΔTP_DM3^Col-0^ and mutant (*; p < 0.05). (L) Expression of DM3 mutants with DM2h^Bla-1^ in *Nb*. DM2h^Bla-1^ was driven by endogenous promoter, while _ΔTP_DM3s and GFP were driven by 35S promoter. Photos were taken five days after infiltration. Numbers indicate the incidents of confluent HR out of two independent experiments with a total eighteen leaves. The scale bar equals 2 cm. (M) Western blot of DM3 mutants. *Nb* leaf samples were collected at 32 hours after infiltration. See also Figure S4 to S6.

Hydrophobicity of isoleucine at 165 in _ΔTP_DM3^Hh-0^ results in the loss of interaction among T165, S167 and D332 observed in _ΔTP_DM3^Col-0^, leading to a melting of the DH in _ΔTP_DM3^Hh-0^. Consequently, M169 and L170 are flipped out and lose the interaction with F425. Furthermore, Q171 in _ΔTP_DM3^Hh-0^, which stabilizes the DL in _ΔTP_DM3^Col-0^, is swung away, abolishing the interaction between Q171 and the backbone of DL (Figures 4C and 4D). Disruption of the secondary structure of DH by the T165I polymorphism in _ΔTP_DM3^Hh-0^ concomitantly affects the interaction at the dimer interface in which a part of DL (335-340) becomes totally disordered (Figure 4D). However, A359 has no disruptive effect on the dimer interface of DM3^Hh-0^, in which interaction between D358 and N349 is retained (Figure S5).

As structural analysis of _ΔTP_DM3^Col-0^ and _ΔTP_DM3^Hh-0^ showed disruption of the dimer interface in _ΔTP_DM3^Hh-0^, we reasoned that the dimer interface disturbance in DM3^Hh-0^ may influence the stability of the complex. Therefore, we performed a thermal shift assay (TSA) and measured the melting temperature (T_m_) of _ΔTP_DM3^Col-0^ and _ΔTP_DM3^Hh-0^ proteins. T_m_ of _ΔTP_DM3^Hh-0^ was measured as 63.0 °C, substantially lower than that of _ΔTP_DM3^Col-0^ (71.5 °C) (Figure 4E), indicating that the polymorphisms found in DM3^Hh-0^ destabilize the DM3 complex. We further performed the prolyl aminopeptidase (PAP) assay to examine whether the polymorphism affects the catalytic activity of DM3. Despite the decreased stability, the PAP activity of _ΔTP_DM3^Col-0^ and _ΔTP_DM3^Hh-0^ proteins were comparable (Figure 4F). To functionally validate the observed structural and biochemical changes *in planta*, we performed an agrobacterium infiltration assay in *Nb*, co-expressing DM2h^Bla-1^ with _ΔTP_DM3^Col-0^ containing either T165I or T359A polymorphisms (_ΔTP_DM3^Col-0^ T165I and _ΔTP_DM3^Col-0^ T359A). Only _ΔTP_DM3^Col-0^ T165I resulted in HR with DM2h^Bla-1^, indicating that T165I-mediated dimer interface disruption at DM3 is critical for DM2h^Bla-1^-mediated immune responses in plants (Figure 4G and 4H).

### Structure-guided dimer interface mutants triggering cell death with DM2h^Bla-1^

The two cryo-EM structures and biochemical properties of DM3 variants suggest that there exists a structural determinant at the dimer interface of DM3 responsible for hybrid necrosis. To verify this idea, we generated a series of structure-guided mutants of _ΔTP_DM3^Col-0^ that would interfere with stabilizing interactions at the dimer interface. In the dimer interface of DM3^Col-0^, D332, Q345, and N349 are located opposite to T165 and T359 (Figure 4I). We mutated those amino acids to arginine or alanine to induce steric hindrance or abolish electrostatic interaction, respectively, and produced D332R, Q345R, DQ-RR (D332R Q345R double mutant), and DQN-AAA (D332A Q345A N349A triple mutant) mutant proteins of _ΔTP_DM3^Col-0^. We found that all recombinant dimer interface mutants of DM3s were eluted at the DM3 hexamer peak from the gel-filtration chromatography (Figure S6A and S6B). TSA analyses demonstrated that, except for Q345R, all single, double and triple interface mutants had reduced T_m_ similar to that of _ΔTP_DM3^Hh-0^ (63.0 °C) (Figure 4J). Like _ΔTP_DM3^Hh-0^, all dimer interface mutants showed PAP activity (Figure 4K). In the *Nb* assays, mutants harbouring substitutions at D322 in particular appeared to gain HR-triggering capability. Even single substitution D332R rendered _ΔTP_DM3^Col-0^ capable of triggering HR with DM2h^Bla-1^, albeit less consistently than _ΔTP_DM3^Hh-0^, while the triple mutant DQN-AAA showed consistent HR with DM2h^Bla-1^. HR-triggering capability appears to be associated with protein stability *in planta*, as the DQ-RR mutant with the least consistent HR frequency (3/18) showed its expression level much lower than other mutants *in planta*. No HR was observed for Q345R mutant, albeit obvious protein accumulation, consistent with its TSA result comparable to that of _ΔTP_DM3^Col-0^ (Figures 4J, 4L, and 4M). From TSA data and HR assays, we conclude that the destabilization of the dimer interface rather a specific mutation (T165I) is key to DM3’s ability to trigger HR with DM2h^Bla-1^.

### The trimer interface of DM3 is dispensable for immunity

As dimer interface mutants trigger HR with DM2h^Bla-1^, we next tested whether the disruption of the trimer interface, supposedly influencing the maintenance of hexamer architecture of DM3, would affect the NLR-mediated cell death. Based on the cryo-EM structure, we produced trimer interface mutants which would lose tight interactions between dimers, by truncating the last five C-terminal residues (ΔC; Δ511-515; ΔKKPLF) or mutating hydrophobic amino acids on the trimer interface to glutamates, to induce charge repulsion (YV-EE; Y460E V484E) on the _ΔTP_DM3^Col-0^ background (Figure 5A). From gel-filtration, both trimer interface mutants were eluted after the hexamer peak, indicating that the hexamer configuration is not maintained (Figure S6C and S6D). T_m_ of the YV-EE and ΔC mutants was determined as 45.5 °C and 44.0 °C, respectively, much lower than that of _ΔTP_DM3^Col-0^ wild type, _ΔTP_DM3^Hh-0^ and dimer interface mutants (Figure 5B). Additionally, PAP activity of the trimer interface mutants was impaired (Figure 5C). Co-expression of the trimer interface mutants of _ΔTP_DM3^Col-0^ with DM2h^Bla-1^ did not cause HR, suggesting that mere presence of DM3 dimer with the dimeric interface configured as Col-0 form does not suffice the immune activation (Figure 5D and 5E). We further tested the effects of trimer interface disruption using the autoimmunity variant _ΔTP_DM3^Hh-0^ using the same C-terminal truncation. When infiltrated with DM2h^Bla-1^, _ΔTP_DM3^Hh-0^ ΔC still expressed and triggered robust HR (Figures 5H and 5I), suggesting that maintenance of hexametric configuration is not necessary for immune activation. T_m_ of _ΔTP_DM3^Hh-0^ ΔC was 36°C, even lower than that of the Col-0 counterpart with gel-filtration assay resulting in the peak at 16 mL, equivalent to monomer (Figures 5B and S6C-S6D). As expected for a mutant not being able to maintain the hexamer form, PAP activity of _ΔTP_DM3^Hh-0^ ΔC was not detectable (Figure 5C). Taken together, the integrity of the dimeric interface, but neither overall maintenance of hexamer nor the integrity of the trimer interface, is the main determining factor for DM2h-mediated immune activation.

**Figure 5.**
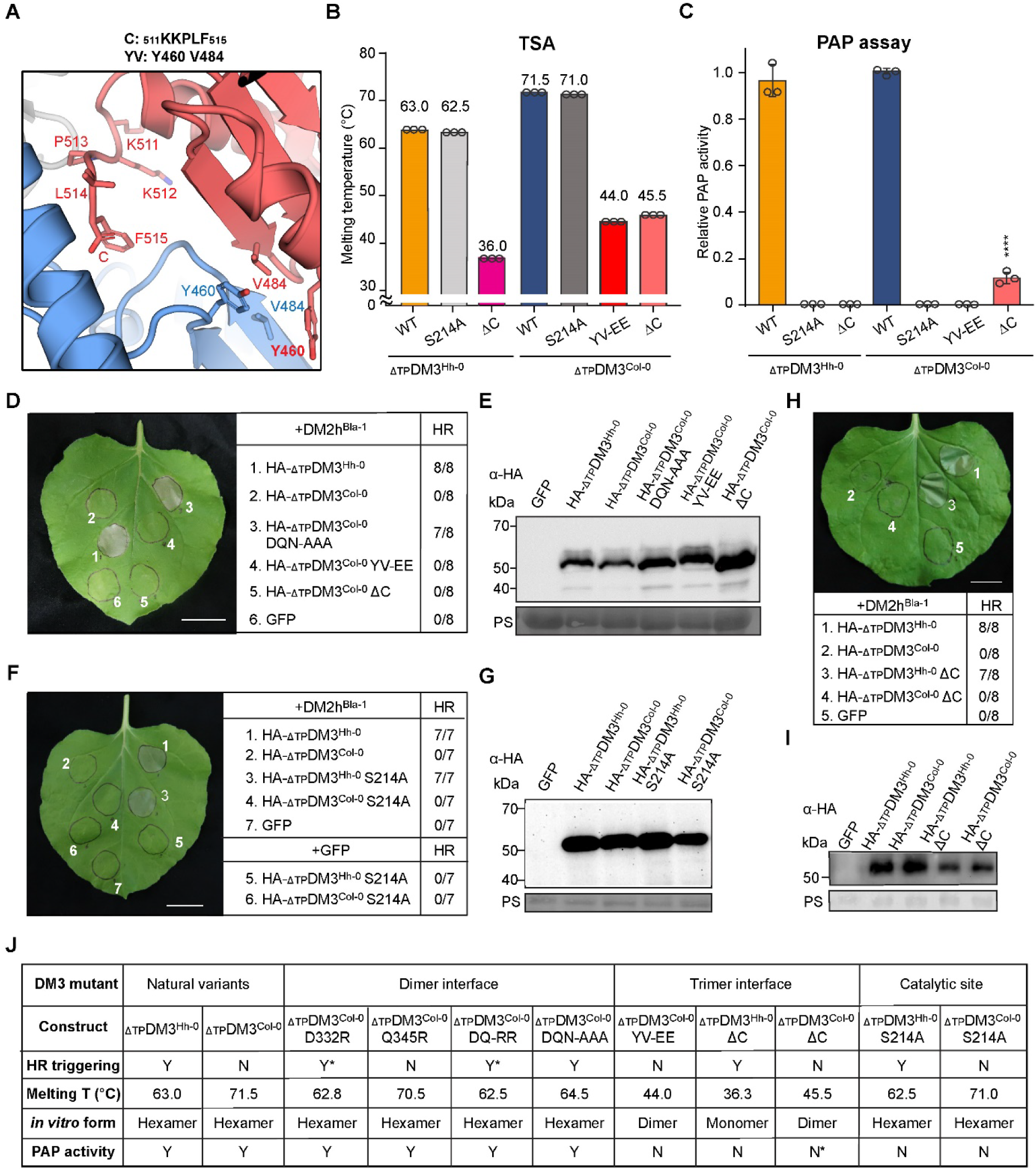
Structure-guided trimer interface mutations and catalytic site mutations impair PAP activity, but not cell-death triggering ability of DM3^Hh-0^. (A) The trimer interface of DM3^Col-0^ shows C-terminal residues (511-515, KKPLF; herein C) and hydrophobic residues (Y460 and V484; herein Y and V). (B) TSA shows the melting temperature of purified trimer interface mutant proteins (ΔC; Δ511-515 and YV-EE; Y460E V484E) and catalytic site mutant proteins (S214A) of _ΔTP_DM3^Col-0^ and _ΔTP_DM3^Hh-0^. Each data point is represented by a circle (n = 3). (C) The enzymatic activity assay shows normalized PAP activity of the trimer interface mutants. Individual data points (n = 3) and standard deviation are represented, and asterisk indicates p-value from two-tailed t-test between wild type _ΔTP_DM3^Col-0^ and mutants (****; p < 0.0001). _ΔTP_DM3^Col-0^ S214A, _ΔTP_DM3^Col-0^ YV-EE and _ΔTP_DM3^Hh-0^ S214A show no PAP activity. (D)(F)(H) Expression of DM3 mutants with DM2h^Bla-1^ in *Nb*. The expression of DM2h^Bla-1^ was driven by endogenous promoter, while _ΔTP_DM3s and GFP were driven by 35S promoter. Photos were taken five days after infiltration. The numbers indicate leaves with fully developed HR out of indicated number of leaves. Scale bar equals 2 cm. (E)(G)(I) Western blot of DM3 mutants. Agroinfiltrated *Nb* leaf samples were collected at 28 hours after infiltration. PS: Ponceau S. (J) Summary of DM3 mutants tested in this study. Y: yes; N: no; Y*: less frequent HR; N*: residual activity. See also Figure S6.

### PAP activity of DM3 is separable from immune function

To precisely discriminate functional relevance between HR-triggering and PAP activity of DM3, we mutated the critical residue of catalytic site S214 to alanine (S214A) in the _ΔTP_DM3^Col-0^ and _ΔTP_DM3^Hh-0^ (Figure S6F). While the S214A mutant still maintains the hexameric structure *in vitro* with little effect on the T_m_ (Figures 5B and S6E), this change in the catalytic core abrogated the PAP activity (Figure 5C). Despite the loss of PAP activity, co-expression of _ΔTP_DM3^Hh-0^ S214A and DM2h^Bla-1^ in *Nb* still triggered consistent HR as severe as _ΔTP_DM3^Hh-0^ does (Figures 5F-5G), indicating that PAP activity is not required for DM3-activated immunity with DM2h^Bla-1^. These data with PAP activities of other DM3 mutants (Figures 4F, 4K, and 5C) suggest that the PAP activity of DM3 rather correlates with the integrity of hexameric complex but not with HR-triggering capabilities of DM3 variants (Figure 5J).

### Instability of DM3^Hh-0^ hexamer associated with immunity in plants

We next examined the oligomeric status of DM3 variants *in planta* to address the biophysical status of DM3 responsible for activating DM2h NLR. First, we attempted to verify protein-protein interaction between DM3 molecules using bimolecular fluorescence complementation (BiFC) in the *Nb* system that we used for assaying the immune responses. Fully recovered fluorescent YFP signal was observed throughout the cytoplasm of pavement cells when the non-fluorescent N-terminal fragment of YFP (nYFP-:1-172) fused to the N-terminal end of _ΔTP_DM3^Col-0^ was co-infiltrated with the non-fluorescent C-terminal fragment of YFP (-cYFP:172-238) fused to the C-terminal end of _ΔTP_DM3^Col-0^ (Figure 6A). When the two non-fluorescent tags were attached to the same positions of _ΔTP_DM3^Hh-0^, the BiFC signal was reduced but detectable as compared to the negative controls expressing each of the individual constructs with an empty vector (Figure 6A). Next, we tested *in planta* self-association of DM3 proteins using the *Nb* leaves co-infiltrated with differently tagged _ΔTP_DM3, either with HA or FLAG at its N-terminus. We performed immunoprecipitation (IP) using anti-FLAG agarose beads, which successfully enriched both variants of DM3. The HA-tagged _ΔTP_DM3^Col-0^ was readily detectable within the IP fraction containing the Col-0 variant, while the detection of HA-_ΔTP_DM3^Hh-0^ was relatively weak in the IP fraction containing the Hh-0 variant compared with that of the Col-0 counterpart (Figure 6B). Both *in planta* BiFC and co-immunoprecipitation (co-IP) data were consistent with the biophysical properties we assayed with recombinant proteins, corroborating our findings on both DM3 variants self-associating but with different affinity.

**Figure 6.**
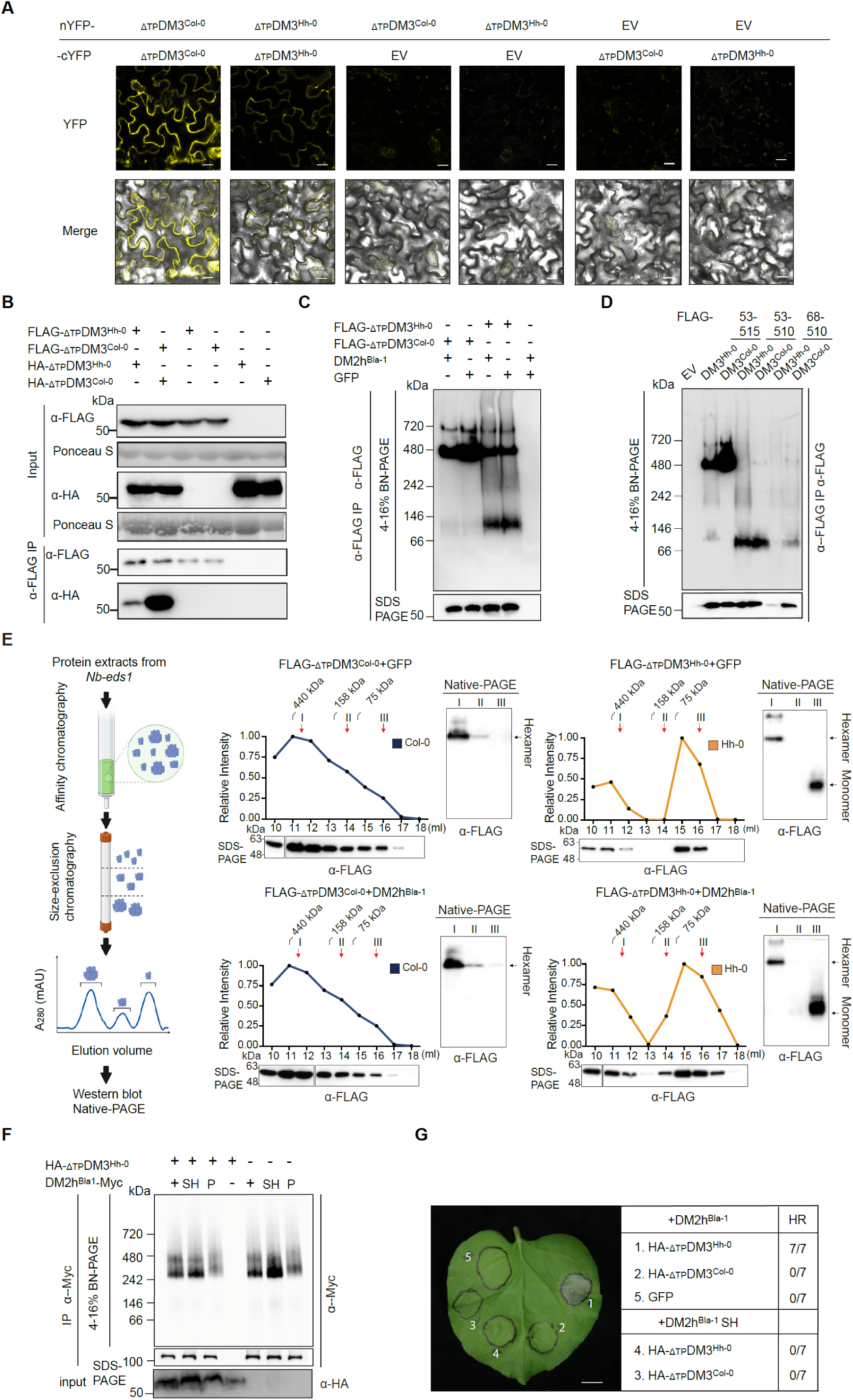
*In planta* oligomer assessments demonstrate a tight DM3^Col-0^ hexamer and above, and a loosened maintenance of DM3^Hh-0^ hexamer with a leakage of monomers. (A) Self-interaction of DM3 was shown by the BiFC assay. _ΔTP_DM3-cYFP, nYFP-_ΔTP_DM3 or empty vector were co-expressed in *Nb* leaves for detecting YFP signals under confocal microscopy. Photo were taken 30 hpi. Scale bar equals 20 µm. (B) Both DM3^Hh-0^ and DM3^Col-0^ self-interacted *in vivo*. *Nb* leaves infiltrated with the indicated DM3 constructs in combination were used for IP with anti-FLAG Affinity gel. Western blots were performed with anti-HA or anti-FLAG antibody for input proteins and IPed DM3. (C) BN-PAGE gel image was shown to visualize different migrating patterns of IPed DM3 complexes with or without DM2h^Bla-1^ co-expressed in *Nb-eds1*. (D) BN-PAGE gel showing complex formation of IPed DM3 with different length expressed in *Nb*. IPed DM3 was also run in SDS-PAGE as loading control. (E) DM3 purification from *Nb*-*eds1* with schematic illustration of experimental procedures (left) and I, II, III indicate hexamer, dimer and monomer sizes, respectively. (F) IP followed by BN-PAGE shows DM2h-Myc variants detected as complexes both in resting state (co-expressed with GFP) and activated state (with DM3^Hh-0^). SH refers to SH to AA mutations at the AE interface in TIR domain; P: p-loop mutation (GI<u>GK</u>TT^299-304^ to GI<u>AA</u>TT). Infiltrated *Nb eds1* leaves were collected 40 hpi for all the experiments shown in this figure. (G) Expression of DM2hBla-1/DM2hBla-1 SH mutant with _ΔTP_DM3^Hh-0^/_ΔTP_DM3^Col-0^/GFP in *Nb.* The numbers indicate leaves with fully developed HR out of indicated number of leaves. Scale bar equals 1 cm. See also Figure S6.

To further examine the oligomerization status of DM3 proteins *in vivo*, we performed blue native-polyacrylamide gel electrophoresis (BN-PAGE) using the immunoprecipitated (IPed) DM3 proteins expressed in *Nb-eds1* to avoid potential protein degradation associated with cell death. The majority of _ΔTP_DM3^Col-0^ migrated in BN-PAGE at a size ∼480 kDa with an additional slowly migrating band detectable at ∼720 kDa along with faint traces faster migrating below the size marker of 146 kDa (Figure 6C). The migration pattern of _ΔTP_DM3^Hh-0^ in BN-PAGE was different from that of _ΔTP_DM3^Col-0^ such that the distribution of the three migrating forms were shifted down to faster migrating forms. The proportion of the major form around 480 kDa was reduced with the obvious appearance of the faster migrating one below 146 kDa (Figure 6C). To associate the major bands with oligomeric status, we evaluated the migration pattern of recombinant _ΔTP_DM3^Col-0^ proteins in BN-PAGE, confirming that the hexamer migrated with a major form around 480 kDa along with a minor form around 720 kDa. Boiled sample, supposedly reducing the oligomeric status, migrated below 146 kDa with a trace around 242 kDa that was only visible in western blot but not in Coomassie blue staining (Figure S6I). Thus, we concluded that the NLR-activating form of DM3^Hh-0^ is less proficient in maintaining a tight hexameric or even higher oligomeric status in the plant cell, rendering their units be prone to disassemble from the oligomers. To further explore oligomeric dynamics of DM3 *in vivo*, we tested *in planta* expressed _ΔTP_DM3 ΔC (53-510) mutants that are defective in assembling dimers into the trimer of dimer configuration in BN-PAGE, of which recombinant proteins came out as dimer or monomer in gel filtration assays (Figure S6C). We also included a further reduced fragment of DM3 (DM3 68-510), which we defined as the smallest unit capable of triggering DM2h-mediated HR in *Nb* and of maintaining structural integrity (Figures S6G and S2). These trimer-interface defective forms of DM3 migrated largely as monomeric forms with a small trace of hexamer and faint traces of dimeric forms visible below the marker sized at 242 kDa (Figure 6D). Additionally, DM3^Hh-0^ fragments were less detectable than those of DM3^Col-0^ such that DM3^Hh-0^ 68-510 was barely detectable *in planta* despite its HR-triggering ability (Figure S6H), suggesting that DM3 protein instability is associated with disassembly from the oligomeric units. Importantly, the detection of monomeric pool of DM3^Col-0^ *in planta* clearly indicates that the mere presence of monomer is not sufficient for DM2h NLR activation, and the loosened dimeric interface as configured as Hh-0 form is the critical feature switching on NLR activation

Next, we scaled up the expression of _ΔTP_DM3 proteins in *Nb-eds1* and size-separated the IPed DM3 protein pools using size exclusion chromatography (Figure 6E). We first correlated the standards for gel filtration with the recombinant DM3 variants of which oligomeric status is known, verifying that hexamer, dimer and monomer run around the standards of 440 kDa (I), below of 158 kDa (II), and below of 75 (III), respectively (Figure 6E, red arrows). The gel filtration profile of DM3^Col-0^ from plants showed a gradual decrease peaking at a fraction right above the size marker 440 kDa irrespective of the presence or absence of DM2h, indicating the DM3 complex formation is not affected by its partner NLR. Subsequent running of the fractions corresponding to hexamer (I), dimer (II) and monomer (III) in 7% Tris-Glycine native-PAGE showed that the fraction I migrated as a major band with a trace of upper band, consistent with the BN-PAGE result. Despite the lower size matching with dimer and monomer, the fraction II and III still migrated in the native gel as hexamer without any detectable traces below the size of hexamer, indicating that *in planta* DM3^Col-0^ pool preferentially maintain higher oligomeric status as hexamer and above. We found that the size separation profile of DM3^Hh-0^ pool strikingly differs from that of DM3^Col-0^. There was a noticeable reduction in the protein amount in the fractions spanning dimeric configuration, which divided the gel filtration profile into two major peaks instead of a continuously tailing pattern seen in that of Col-0 (Figure 6E). Native gel analysis revealed that there is a clear presence of fast-migrating DM3^Hh-0^ pool in the monomeric fraction III, while the migration pattern of hexameric fraction I was equivalent to that of DM3^Col-0^. Fraction II of samples expressing DM3 alone with GFP control or with DM2h^Bla-1^was barely detectable in the native gel compared to the other fractions, presumably reflecting the fleeting nature of dimer.

Next, we examined whether DM2h^Bla-1^ NLR oligomeric status is affected by the presence of hybrid necrosis triggering DM3^Hh-0^. When IPed DM2h NLR pools were run on a BN-PAGE gel, there exist two discrete migrating forms of DM2h, of which oligomeric status can be inferred as dimer and tetramer based on their sizes between 242∼480 kDa (Figure 6F). While mutations in P-loop (from GI<u>GK</u>TT^299–304^ to GI<u>AA</u>TT) in the NLR rendered the bands less discrete, substitutions of S111 and H112 to alanine (SH-AA) in the AE interface critical for assembling TIR-domains as a dimeric unit^47,48^ was not sufficient to break the dimer or tetramer to monomeric units. Rather, the dimeric band of DM2h^Bla-1^ (SH-AA) in the absence of DM3^Hh-0^ was more conspicuous than non-mutated DM2h^Bla-1^, suggesting the possibility of DM2h^Bla-1^ dimers engaging another interface for maintaining dimer status in resting state. As we observed no noticeable major differences in the NLR configuration in the absence and presence of DM3 trigger, we conclude that DM3 controls NLR activity not through inducing higher oligomeric NLR status *per se* but presumably by inducing subtle changes in the preassembled NLR units to structurally reconfigure the catalytic TIR centres. Although no oligomeric status changes were observed, mutations at SH made DM2h^Bla-1^ non-functional protein as it failed to trigger HR with DM3^Hh-0^ in *Nb* (Figure 6G), supporting the importance of AE interface in the active status as well as the notion of reconfiguration within the preassembled NLR units. Despite successful enrichment of DM2h with IP experiments (Figure 6F), we failed to detect DM3 variants within the IPed pool (data not shown), suggesting that the activation of DM2h by DM3 relies on transient nature of physical interaction or unknown indirect mechanism.

## DISCUSSION

Using the two-component autoimmune system discovered from hybrid necrosis studies, we have presented an atomic resolution of the cause of genetic incompatibility that inadvertently activates host immunity in plants. Our cryo-EM data highlights key structural determinants in the ordered complex of DM3 responsible for activating the cognate NLR, which lies in its tight packing at the dimer interface rather than its catalytic peptidase activity. Having a fine-scale understanding of host enzyme packing and its regulation sheds light on potential host cellular events that can instigate NLR-mediated immune responses. Given the fact that DM2h NLR used in this study is encoded by one of the most rampant multigene NLR clusters in the genome^49–51^, it is likely that our current study highlights one of many uncountable guarding strategies that NLRs employ to oversee cellular states.

### Orderly assemblage of DM3 into high-order structures enables peptidase and immune function distinctively

Our data shows that the immune function of DM3 is independent of its catalytic function as a peptidase, and this distinction is conditioned by its orderly quaternary structure. As revealed by cryo-EM structure determination and *in planta* assays, the majority of DM3 is likely to assemble as a tight hexamer in the cell, which we determined is an essential packing unit for the peptidase activity (Figure 5). Previously, DM3/PAP2 has been proposed to function in the mitochondria to process peptides with proline at its N-terminal end^42,43^, although this notion does not exclude the role of PAP activities in regulating cellular proline homeostasis. Such hexamer packing has been reported for Tricorn protease and Lon protease, in which catalytic cores are also buried within the barrel as seen in the DM3 hexamer^52–57^. Thus, it appears that the maintenance of such hexamer or even further packing as a higher unit potentially serve as a regulatory checkpoint integrated into the orderly assembly of these degrading entities in the cell. Consistent with this, we found that DM3 tends to form a higher oligomeric status *in vivo* and *in vitro*. In the *Nb* expression system, a slowly migrating band above the hexamer was detected (Figures 6C-6E). We also found an *in vitro* assembly of a filament structure of DM3^Col-0^ under a specific condition as shown by negative staining EM (Figure S7). Recent studies with cryo-EM determination uncovered that various enzymes can be assembled as ordered filaments in a helical packing, which often serves to regulate catalytic activities upon environmental or physiological changes^58–61^ Thus, it will be interesting to test if the filamentous status of DM3 serves a similar regulatory role for enzyme activity and/or immune function such as to stabilize the dimer interface.

For the case of DM3, we found that the majority of naturally existing forms would assemble as a tight hexamer as exemplified with the Col-0 variant, in which the dimer- and trimer interfaces enable ordered packing of the molecules. It is known that proteins with ABH domains feature structural flexibility as they can easily accommodate additional protein domains. Our structural determination also shows that the DL located in the lid domain and DH inserted within the ABH domain of the DM3 monomer are key regions for DM3 dimerization. It is notable that the T165I polymorphism on the rare variant DM3^Hh-0^ poised for immune function affects neither the hexamer conformation nor PAP activity, indicating that the immune function of DM3 is separable from PAP function. We rather confirmed that the polymorphism leading to the structural disruption in DH and to disordering DL at the very local area in the dimer interface reduces thermostability of the whole hexamer and eventually compromises the orderly assembly of oligomers. Our structure-function studies of the two DM3 complexes pinpointing an immune-regulatory role embedded in the dimer interface is somewhat analogous to the EP domain interface between EDS1 and PAD4, which confer their immune function without a contribution from the catalytic cores^62^. While the structural superimposition failed to align the EDS1-like ABH proteins with DM3 due to flexible juxtaposition of the lids over the ABH domains, it remains to be further explored whether the instability-inducing structural disruption found in the DM3^Hh-0^ dimer can provide any pockets for potential second messenger molecules.

### Structural disruption at the DM3 dimer interface activates DM2h NLR

Although *in vitro* and *in planta* assessments of oligomeric status confirmed that both DM3 variants maintain hexamer status albeit with different stability, we found that the maintenance of hexamer is not necessary for the DM3 immune function (Figures 5D, 5H-5I, and 6D). Furthermore, we uncovered that it is not the production of monomeric DM3 pool itself that is required for the NLR activation but a disordered or altered structure at DH/DL area of dimer interface present in whichever pool of DM3 oligomers. This notion is evidenced by the data of all the trimeric interface mutants of DM3^Col-0^ that we tested, falling out as monomers in plants, failed to activate DM2h NLR, while equivalent fragments of DM3^Hh-0^ were sufficient in triggering immunity even if they were barely detectable presumably due to instability associated with disassembly of oligomers (Figure 6D).

A likely scenario of host cellular events that activate the TIR-catalytic activity of DM2h NLR would start with an induced DM3 instability specifically at the surface-exposed dimeric interface surrounding DL and DH, which can be potentially triggered by pathogens or cellular stresses. This instability can be readily followed by partial disassembly of the trimer of dimers or an altered equilibrium of monomer, dimer and hexamer status. Given the orderly units in the hexamer and rarity of detectable dimer pools in plants (Figure 6E), we speculate that DM3 dimer units falling out from the packing unit with intrinsic instability surrounding DL/DH instantaneously disperse as monomers. Our favoured hypothesis regarding the immune switch is that an exposure of the DH/DL interface either from an ephemeral dimer unit or even from hexamer is the key structural event that DM2h NLR monitors as a proxy of cellular homeostasis. In other words, the event can induce a reconfiguration of the preassembled NLR dimer or tetramer into a proper tetramer unit as a holoenzyme with strong NADase catalytic activity. Whether or not this gaining of TIR-catalytic activity of DM2h NLR is mediated by direct interaction between the dimer units of DM2h and DM3 remains to be further investigated with a technology that is capable of capturing transient interactions. Unfortunately, we did not succeed in detecting the physical interaction *in vivo* and *in vitro* using co-IP experiments. However, we also do not exclude the possibility of DM3 indirectly activating DM2h NLRs by other means, such as small molecule sequestration.

### DM2 NLRs monitor proteins with distinct functions related to cellular homeostasis

It is important to note that the *DM2/RPP1* locus is a hot spot spawning hybrid necrosis alleles in Arabidopsis, with at least five different genetic partners known to date^63,64^. Best studied examples include natural alleles of *DM2h* that are prone to trigger autoimmunity not only in hybrid backgrounds but also by chemically induced mutations in the same genetic background. Such host proteins that DM2hs make surveillance on include the receptor kinase SRF3 that coordinates iron homeostasis^37,38^, the cysteine synthase O-acetylserine (thiol) lyase (OASTL) involved in maintaining organic cysteine and sulfur levels^39,65^, DM3 which supposedly regulates proline homeostasis and stress responses^4,42^, and intriguingly nuclear pool of EDS1^5,41^. While most of our knowledge on NLR activation comes from studies of effector-triggered immunity, DM2h NLRs participating in such diverse autoimmune events make it a unique candidate to study immune activation by cellular stresses.

In this work, we demonstrated that the *in vitro* PAP catalytic activity directly associated with proline production is non-differentiated between DM3^Col-0^ and DM3^Hh-0^. Nonetheless, we also found that relative abundance of hexamer, a structural requisite for the PAP activity, is different between the two variants in the plant cell. It is likely that DM3’s PAP activity is compromised to some extent when the complex adopts the form like DM3^Hh-0^ due to the reduced hexamer stability. Without an accurate assessment of proline amount in the plant cell, it is premature to propose that the NLR activation could be triggered by homeotic disturbance of actual proline level in an organism with a compromised DM3 hexamer configuration. Although we currently prefer the hypothesis of DM3 inducing structural conformation changes to the preassembled NLR oligomers through direct or indirect interactions, we cannot exclude the possibility that the hexamer instability can be seen by DM2h NLR as a proxy of proline level changes. Indeed, proline is considered as a stress indicator and regulator in plants^66^, and *DM3* loss-of-function mutants show hypersensitivity to various abiotic stresses^42^. Given that DM2h NLRs monitor the sanity of various enzymes regulating the homeostasis of iron, cysteine and proline, a tantalizing hypothesis arises for the mode of activation of DM2h NLRs involving recognition of cellular stress levels. Whether this mode of NLR recognition directly involves actual molecules, such as accumulation of peptides or depletion of free amino acids, or indirect sensing of a physiological phenomenon shared among these cellular stresses remains to be elucidated. Guarding multiple related molecular processes are known for animal NLR, NLRP3 as a strategy to recognize cellular stresses to initiate inflammasome assembly^67,68^. It remains to be seen how very closely related NLRs and likewise distantly related ones adopt similar strategies to monitor various cellular status through guarding host enzymes and molecules.

In this sense, our finding of DM2h^Bla-1^ NLR remaining as preassembled dimers or even tetramers in the plant cell in the absence of DM3^Hh-0^ is intriguing. The preassembly of autoimmune-risk NLRs were previously reported for DM1 and DM2d, which are TIR-containing NLRs, and DM6 NLR that contains CC-domain at its N-terminus^64,69^. Preassembled dimers of NRC2 helper NLR were recently shown to adopt different structural interface than those found in the active hexamer of NRC2, suggesting that there has to be a rearrangement of the interfaces during NLR activation^70^. Breaking the preassembled interface of NRC2 dimer increased their sensitivity to sensor NLRs and even led to autoactivity, indicating that subtle changes in the dimeric configuration can place NLR molecules in incremental steps toward activation. We speculate that our DM NLRs residing in a multigene NLR cluster gained an elevated status in the incremental activation scheme, as proposed by domain swap studies of NLRs^4,71,72^, so that the preassembled dimers function as a safeguard not to overly activate unless triggered, while this autoinhibition is labile and sensitized to cellular stresses. It will be interesting to perform comparative studies of closely related DM2 NLR alleles and chimeras in terms of assembling oligomers upon imposed cellular stresses. Addressing the structural features of resting state of NLRs will also be instrumental to understand how these labile switches operate in the cell.

### DM2h guarding DM3 for immunity?

Based on the biochemical studies of guardee/decoy modification by effectors, it has long been proposed that conformational changes of a host molecule under an NLR surveillance could serve as an immunity activation cue. Several biochemical examples of recognition modes were uncovered for guardee/decoy, such as a disappearance or post-translational modification of the host molecules in the cell serving as cues to NLR activation. However, structural studies on how these biochemical changes translate to conformational changes have lagged behind. Our results demonstrated that the TP is dispensable to activate immunity *in planta*, suggesting a cytoplasmic pool of DM3 is likely being monitored by DM2h. When the pool is maintained as tight hexamers, presumably contributed by DH/DL-assisted packing at the dimer interfaces as the Col-0 form, DM2h NLR does not activate. As the major structural difference between DM3^Col-0^ and DM3^Hh-0^ is located at the hexamer surface, it raises a possibility that a certain conformational change surrounding this region serves as the point of immune surveillance by DM2h NLR. It remains to be determined whether DM3 serves as an effector target, and if so whether the effector-targeting can induce such specific hexamer instability seen in DM3^Hh-0^. Although we lack strong evidence for DM3 as an effector target at the moment as our discovery stems from autoimmunity research, we note that OASTL, another enzyme that DM2h monitors, has been discovered as an effector target from large-scale screens^73^. In addition, we found that the critical change such as T165I introducing bulky side chain at the DM3 dimer interface can be potentially introduced by post-translation modifications such as acetylation, as there were studies using the T to I substitution as an acetylation mimic of threonine^74,75^. Given that several effectors were shown to carry acetyltransferase activities^74–82^, it will be a fascinating research avenue to define immune function of DM3 in line with effector-triggered immunity.

Considering escalated arms race between effectors and host defence strategies, it is conceivable that a certain virulent strategy against current effective NLR-triggered immunity might have already gone extinct^83^. To this end, leveraging an NLR-triggered autoimmunity by host factors provides us with an excellent probe to uncover host cellular processes potentially important for ETI. Here, we showcase its utility by uncovering structural determinants in an enzyme complex. Further investigation of *DM* autoimmunity cases is likely to shed light on host biology which NLRs monitor for the sanity check of cellular homeostasis.

## ACKNOWLEDGMENTS

We thank Siaowan Fong and Persis Chan for their technical assistance. We are grateful to the KARA (KAIST Analysis center for Research Advancement) for management of the negative stain EM and Glacios cryo-TEM. We thank the IBS (Institute for Basic Science) and GSDC (Global Science Experimental Data hub Center) for the cryo-EM data collection. We thank Dr. Johannes Stuttmann for giving *Nb-eds1* and *Nb-nrg1* seeds as gifts. The study was supported by Ministry of Education, Singapore under its Academic Research Fund (MOE-T2EP30221-0012), Singapore National Research Foundation under its Competitive Research Programme (NRF-CRP22-2019-0001), National Research Foundation of Korea (RS-2024-00333346, RS-2024-00344154), KAIST Grand Challenge 30 project (KC30), Global PhD Fellowship (NRF-2018H1A2A1061362).

## AUTHOR CONTRIBUTIONS

Conceptualization: JS, EC

Methodology: GK, WW, NC, NK, YYT, YYL, RRQL

Investigation: GK, WW, NC, NK, YYT, YYL, RRQL, SKN, YZ

Visualization: GK, WW, NC, NK

Funding acquisition: GK, JS, EC

Supervision: JS, EC

Writing – original draft: GK, WW, NC, RRQL, JS, EC

Writing – review & editing: GK, WW, NC, NK, YYT, YYL, RRQL, SKN, YZ, JS, EC

## DECLARATION OF INTERESTS

J.S. is a co-founder and CTO of Epinogen.

N.C. is an employee of Merck Pte Ltd.

## SUPPLEMENTAL INFORMATION

Figures S1–S7

Table S1-S2

**Figure S1. Protein Sequence alignment and the secondary structure of DM3 homologs. Related to Figure 1.**

The alignment portrays conservation between DM3^Col-0^ homologs with indicated important residues. T165 and T359 are labelled with blue triangles to display important DM3 variant residues. Predicted catalytic triad residues S214, D462, and H490 are labelled with red triangles. Based on the DM3 structure in this study, the green star beneath indicates the important dimer interface residues: D332, Q345, and N349; the pink star indicates the trimer interface residues Y460, V484, and C-terminal K511-F515 of the DM3 hexameric complex. The blue box below indicates the N-terminal deletion of DM3^Col-0^ (M1-M51) designed for structural studies. The magenta box below indicates the lid domain region. The protein sequence alignment was done using homology extension (PSI-Coffee) on the T-Coffee server.

**Figure S2. Cryo-EM image processing on _ΔTP_DM3^Col-0^ data and validation on EM map reconstruction. Related to Figure 2.**

(A) A flowchart of the _ΔTP_ DM3^Col-0^ cryo-EM data processing is shown in detail. A representative cryo-EM micrograph, 2D class average, 3D classes and the final EM map (bottom right) are shown. Detailed descriptions of the data processing are in STAR Methods. (B) A FSC curve shows the resolution of the cryo-EM reconstruction of _ΔTP_DM3^Col-0^ as 3.39 Å with 0.143 criteria. (C) Local resolution of the cryo-EM map of _ΔTP_DM3^Col-0^ is estimated and colored by Relion. The scale bar is shown at the bottom. (D) The cryo-EM structure of _ΔTP_DM3^Col-0^ monomer is fitted into the cryo-EM map. The structure of _ΔTP_DM3^Col-0^ monomer and the density are colored dark grey and grey respectively. (E) A structure near the DH (164-172) is fitted in the cryo-EM. T165 is shown with the DH (166-171). (F) A ɑ-helix of lid domain (344-355) is fitted in the cryo-EM density. (G) A part of the Dimerizing Loop (332-342) is fitted in the cryo-EM density. (H) Two β-strands of the ABH domain (79-88 and 102-111) are fitted in the cryo-EM density. (I) A C-terminus of the ABH domain (509-515) is fitted in the cryo-EM density.

**Figure S3. Topology diagram of _ΔTP_DM3^Col-0^ monomer and comparison of individual monomer subunits of _ΔTP_DM3^Col-0^ hexamer. Related to Figure 2.**

(A) The cryo-EM structure of _ΔTP_DM3^Col-0^ monomer is shown. The secondary structure of the lid domain is colored in pale purple. ɑ-helix and β-strand of the ABH domain are colored blue and yellow respectively. (B) The topology diagram of _ΔTP_DM3^Col-0^ monomer shows arrangement of the secondary structures. The secondary structure of the lid domain is labelled with apostrophe (‘). Positions of catalytic triad (S214, D462 and H490) amino acids and the polymorphic residues (T165 and T359) are denoted as black dots. DH, DL and helix insertion are labelled accordingly. (C) Six _ΔTP_DM3^Col-0^ monomers are superimposed and shown as a cartoon. RMSD values between _ΔTP_DM3^Col-0^ subunits are less than 0.286 Å.

**Figure S4. Biochemical characterization, cryo-EM image processing and validation of _ΔTP_DM3^Hh-0^. Related to Figure 4.**

(A) A gel-filtration chromatogram of the purified recombinant _ΔTP_DM3^Hh-0^ is shown. (B) 1 μg of the purified _ΔTP_DM3^Hh-0^ protein was examined by SDS-PAGE. (C) A negative-stain EM shows _ΔTP_DM3^Hh-0^ oligomer. An enlarged micrograph is shown with the scale bar (50 nm). (D) A flowchart of the _ΔTP_DM3^Hh-0^ cryo-EM data processing shows a representative micrograph, 2D class averages and a final EM map (bottom right). Detailed descriptions of the data processing is in STAR Methods. (E) FSC curve shows resolution of the EM map of _ΔTP_DM3^Hh-0^ as 2.89 Å with 0.143 criteria. (F) Local resolution of the cryo-EM map of DM3^Hh-0^ is estimated and colored by Relion. The scale bar is depicted at the bottom. (G) Cryo-EM structure of _ΔTP_DM3^Hh-0^ monomer is fitted in the cryo-EM map. The monomer structure and the cryo-EM map are colored dark grey and grey respectively. (H) The melted DH of the dimer interface of DM3^Hh-0^ (164-172) is fitted in the cryo-EM density. I165 is located before the DH (DH; 166-171). (I) An alpha helix of lid domain (344-355) is fitted in the cryo-EM density. (J) DL (331-344 ; 335-340 are disordered) is fitted in the cryo-EM density. The cryo-EM density for residues 335 and 340 was insufficient to model the structure of DM3^Hh-0^. (K) Two β-strands of the ABH domain (79-88 and 102-111) are fitted in the cryo-EM density. (L) A C-terminus of the ABH domain (509-515) is fitted in the cryo-EM density.

**Figure S5. The T359A polymorphism of DM3^Hh-0^ does not affect the structure of the dimer interface. Related to Figure 4.**

(A) The dimer interface of DM3^Col-0^ shows T165 and T359 (navy) and its interacting amino acids. Electrostatic interactions between amino acids are represented as dotted lines. Dimer interface containing T359 is outlined in a box and shown at the bottom. (B) The dimer interface of DM3^Hh-0^ shows I165 and A359 (orange) and with its interacting amino acids. Electrostatic interactions between amino acids are represented as dotted lines. Dimer interface containing A359 is outlined in a box and shown at the bottom.

**Figure S6. Purification of recombinant structure-guided mutant DM3 proteins for biochemical analysis and the smallest unit of DM3. Related to Figure 4, 5, 6.**

(A) A gel-filtration chromatogram of dimer interface mutants (DQ-RR; D332R Q345R and DQN-AAA; D332A Q345A N349A) and wild type _ΔTP_DM3^Col-0^ and _ΔTP_DM3^Hh-0^ are shown. The standard molecular weight marker is labelled on top. (B) An SDS-PAGE verifies purification of recombinant dimer interface mutation of _ΔTP_DM3^Col-0^. 1 µg of purified protein was loaded on each well. (C) A gel-filtration chromatogram of trimer interface mutants (ΔC; Δ511-515 and YV-EE; Y460E V484E) and wild type _ΔTP_DM3^Col-0^ and _ΔTP_DM3^Hh-0^ are shown. (D) An SDS-PAGE verifies the purity of recombinant trimer interface mutants. 1 µg of purified protein was loaded on each well. (E) A gel-filtration chromatogram of _ΔTP_DM3^Col-0^ and _ΔTP_DM3^Hh-0^ catalytic site mutants (S214A) and wild type _ΔTP_DM3^Col-0^ and _ΔTP_DM3^Hh-0^ are shown. (F) An SDS-PAGE verifies the purity of recombinant catalytic site mutants. (G) Expression of FLAG-tagged truncated DM3 with DM2h^Bla-1^ in *Nb*. DM2h^Bla-1^ was driven by endogenous promoter, while DM3s and GFP were driven by 35S promoter. Photos were taken five days after infiltration. The numbers indicate leaves with fully developed HR out of eight leaves. Scale bar equals 2 cm. (H) Western blot of truncated DM3. Expression of all genes were was driven by 35S promoter. Agroinfiltrated *Nb* leaf samples were collected at 30 hours after infiltration. (I) BN-PAGE of _ΔTP_DM3^Col-0^ recombinant protein. DM3 protein was loaded with or without 95°C 5 minutes treatment, 1 µg for Coomassie blue staining and 1 ng for western blot.

## STAR METHODS

## Key resources table

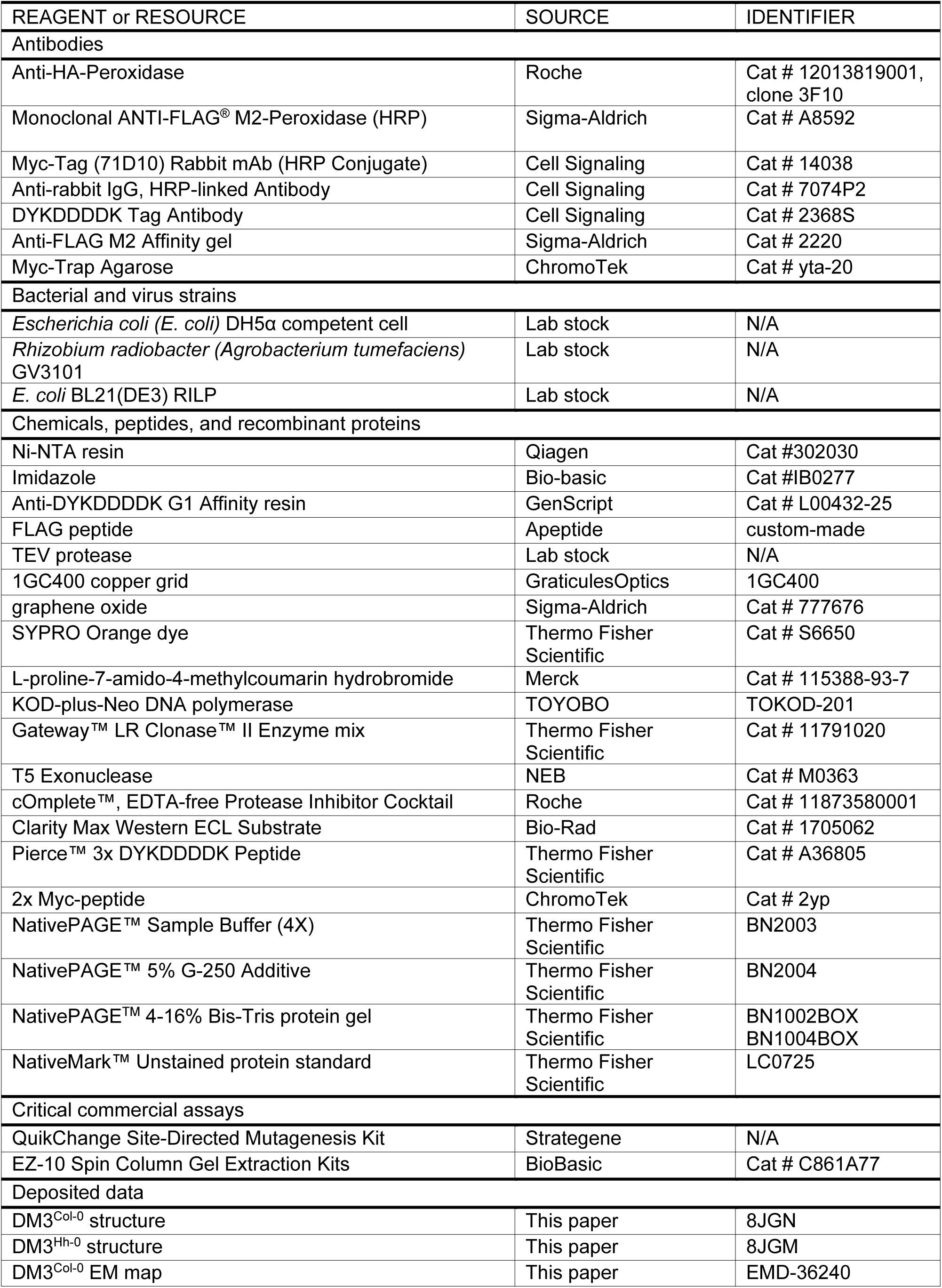
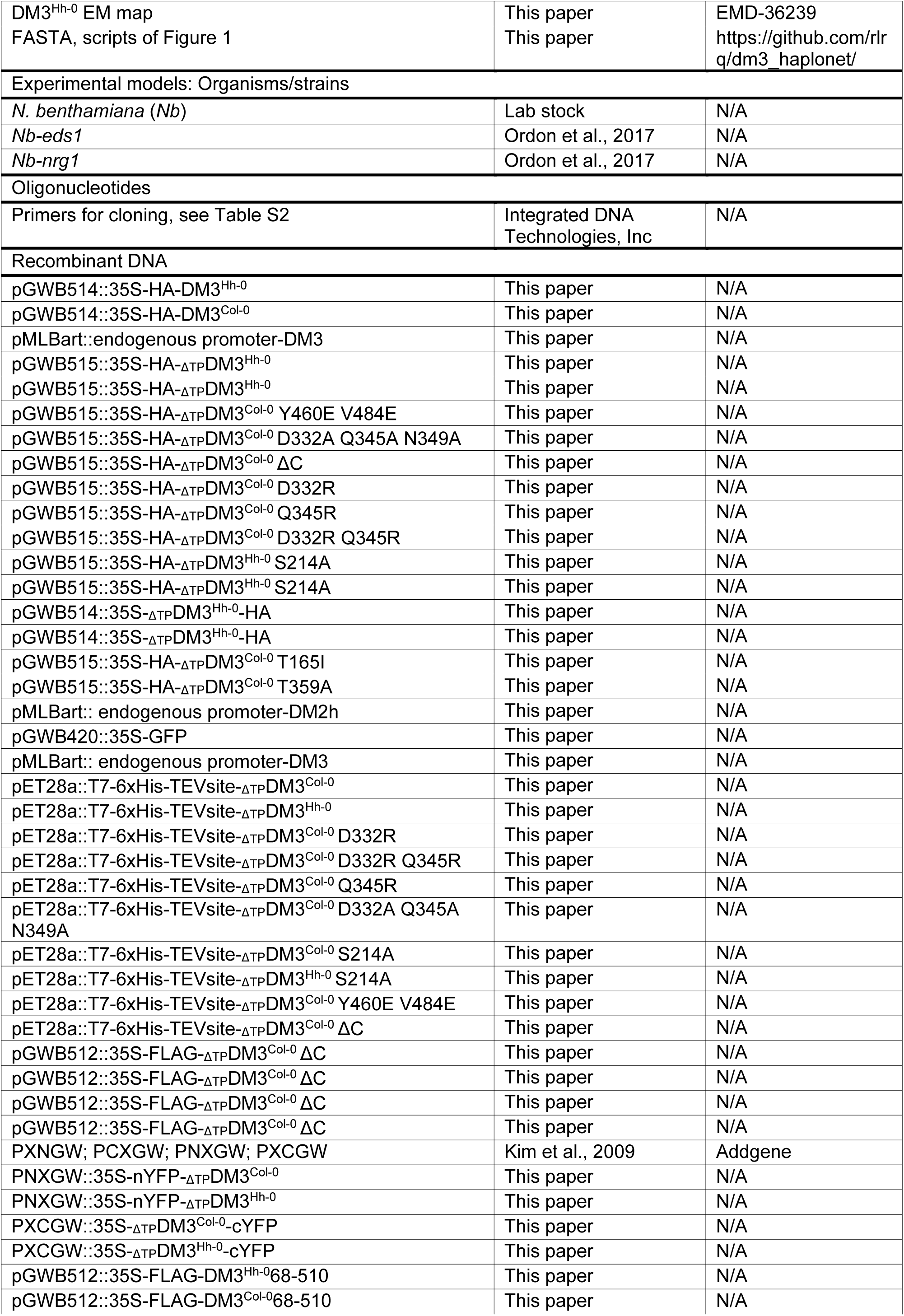
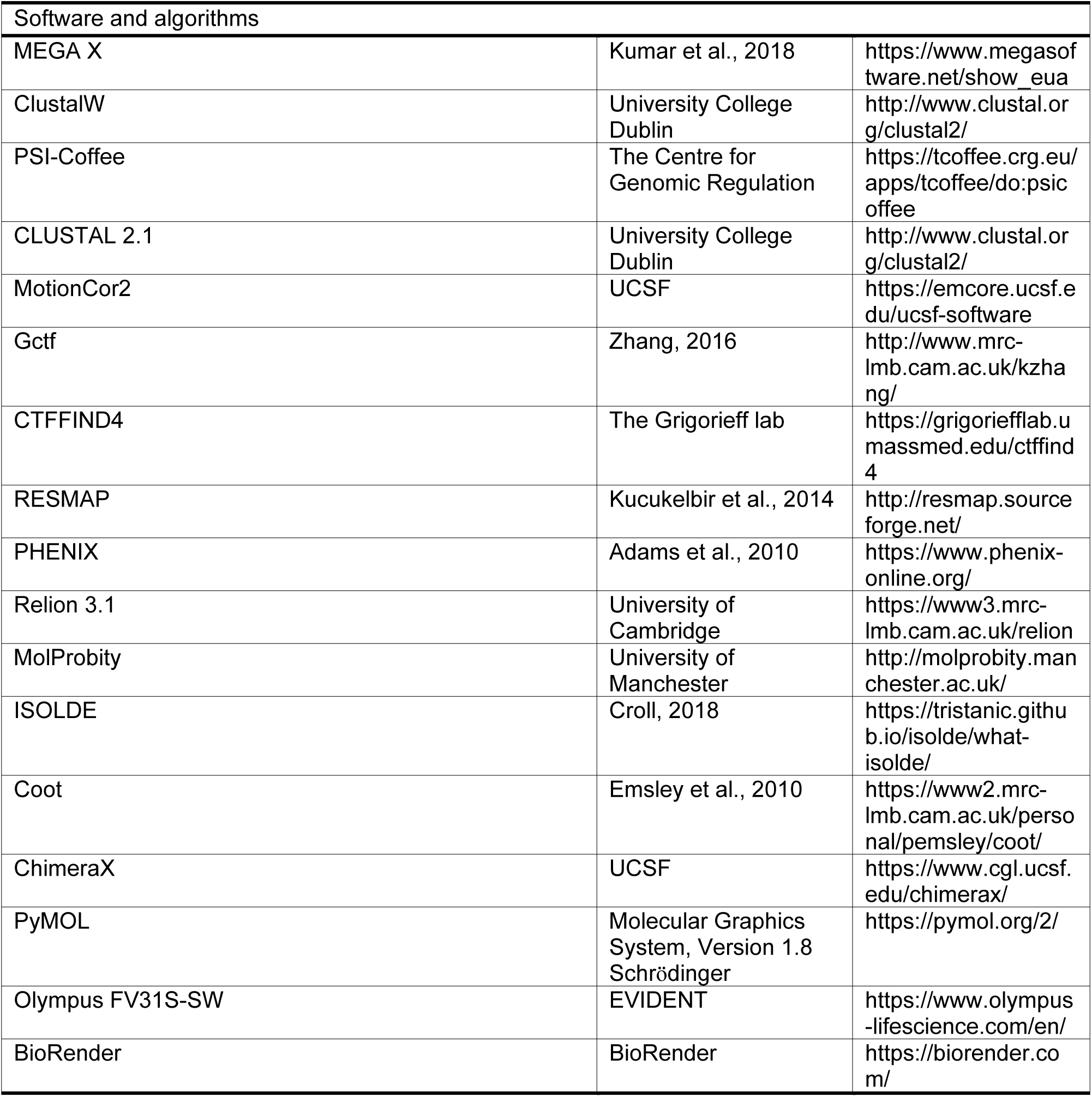

## RESOURCE AVAILABILITY

### Lead contact

Further information and requests for resources and reagents should be directed to and will be fulfilled by the lead contact, Dr. Eunyoung Chae.

### Materials availability

Plasmids generated in this study have been deposited to Addgene This study did not generate new unique reagents

### Data and code availability

For structures of DM3^Col-0^ and DM3^Hh-0^ the atomic coordinates have been deposited in the PDB with accession codes 8JGN and 8JGM, respectively. The EM map for DM3^Col-0^ and DM3^Hh-0^ has been deposited in the Electron Microscopy Database with accession code EMD-36240 and EMD-36239, respectively.

FASTA files containing peptide-as-nucleotide sequences and R script used to generate the haplotype map are deposited at https://github.com/rlrq/dm3_haplonet/.

Any additional information required to reanalyse the data reported in this work paper is available from the lead contact upon request.

## EXPERIMENTAL MODEL AND SUBJECT DETAILS

### Bacteria strain

*Escherichia coli (E. coli)* DH5α

*A. tumefaciens* (strain GV3101)

BL21(DE3) RILP

### Plant materials

*Nb* plants were grown at 24°C in a growth room under a 14 h light/10 h dark cycle. Five-week-old plants were used for *Agrobacterium tumefaciens*-mediated transient expression. *Nb-eds1* and *Nb-nrg1* were gifts from Dr. Johannes Stuttmann.

## METHOD DETAILS

### Sequence analysis

Prediction of transit peptide was done by LOCALIZER^85^. The phylogenetic tree was constructed with MEGA X^86^ by aligning peptide sequences using ClustalW^87^ with default parameters, then generating a maximum-likelihood tree with 1000 bootstrap replicates and all other parameters as default. The protein sequence was aligned using homology extension (PSI-Coffee) ^88^ on T-Coffee server. PSI-Coffee is designed for aligning homologous proteins with no available structural information.

### Haplotype analysis

Coding sequences of DM3 homologues in *A. thaliana* were downloaded using 1001 Genomes^44^ API (application programming interface) and translated. Unknown residues were encoded as Col-0. Resultant sequences were sorted into three bins according to frequency (<10 accessions; 10-100 accessions; >100 accessions) and collapsed. Peptide sequences were re-encoded as arbitrary nucleotides, and these peptide-as-nucleotide sequences were aligned by ClustalW2 from CLUSTAL 2.1^89^. The subsequent alignment was used to generate a haplotype network in R^90^ using pegas version 1.1^91^. FASTA files containing peptide-as-nucleotide sequences and R script used to generate the haplotype map are deposited at https://github.com/rlrq/dm3_haplonet/.

### Cloning, protein expression and purification in *E. coli*

To purify the recombinant DM3 protein, N-terminus truncated construct of DM3 (_ΔTP_DM3; 52-515) was cloned into a modified pET28a with an N-terminal 6xHis-tag followed by a TEV protease cleavage site (ENYLFQ/G). We generated DM3 mutants using the QuikChange Site-Directed Mutagenesis Kit (Strategene). The cloned plasmid was transformed into BL21(DE3) RILP and _ΔTP_DM3 was overexpressed overnight by treating with isopropyl β-D-1-thiogalactopyranoside (IPTG) at 0.6 – 0.8 OD_600_. The cultured cells were collected by centrifugation at 4,000 rpm (4,553 *xg*) for 10 minutes and resuspended with a lysis buffer (50 mM Tris-HCl pH 8.0, 500 mM NaCl, 5% (v/v) glycerol). The cells were lysed by sonication and the lysate was clarified by centrifugation at 13,000 rpm (20,449 *xg*) for 1 hour. The supernatant was then incubated with Ni-NTA resin (Qiagen) for 1 hour and the resins were washed with the lysis buffer supplemented with 20 mM imidazole. _ΔTP_DM3 bound to the resin was eluted with a lysis buffer supplemented with 200 mM imidazole. N-terminus His-tag was removed with TEV protease overnight at room temperature. Cleavage of His-tag was assessed by SDS-PAGE before further purification. The proteins were further purified with a HiTrap Q ion exchange chromatography. The fractions containing _ΔTP_DM3 were then collected and injected to Superdex 200 increase 10/300 column (Cytiva) equilibrated with a gel-filtration buffer (50 mM Tris-HCl pH 8.0, 100 mM NaCl). The recombinant _ΔTP_DM3 proteins were collected and snap-frozen before usage.

### Negative-stain electron microscopy

To observe the purified recombinant _ΔTP_DM3 proteins by negative-stain EM, 3 µL of the protein solution (0.01-0.03 mg/mL) was loaded onto a glow discharged carbon coated 1GC400 copper grid (GraticulesOptics) and incubated for 1 minute. An excess amount of droplet was blotted away and the grid was washed 2 times with distilled water. The grid was then incubated with 1.5 % (v/v) uranyl acetate. After blotting away an excess amount of uranyl acetate, the grid was air-dried before observation. The negative stained grid was observed by the Tecnai F20 transmission electron microscope (FEI) installed at KAIST Analysis Center for Research Advancement (KARA), Korea.

### Single particle cryo-electron microscopy data collection, image processing and refinement

Before cryo-EM sample vitrification, R 1.2/1.3 Cu Quantifoil holey grid (300 mesh) was coated with graphene oxide (GO) based on the GO-PEI method^92^ to resolve a preferred orientation of the _ΔTP_DM3^Col-0^ particles in vitreous ice. 3 µl of the purified DM3^Col-0^ (0.2 mg/mL) was applied onto the GO grid and the grid was blotted for 3 s with -13 blotting force using a Vitrobot Mark IV (Thermo Fisher Scientific). Cryo-EM data were collected by a Glacios cryo-TEM (Thermo Fisher Scientific) with a Falcon III detector installed at KAIST Analysis Center for Research Advancement (KARA). Movie frames were collected in an electron counting mode and each movie frame contains 40 fractions with a total dose of 40 e^-^/Å^2^ at 1.144 Å/pix. The data were processed using Relion 3.1. Beam-induced micrographs motion was corrected with MotionCor2^93^ and contrast transfer function (CTF) of the micrographs was estimated by Gctf ^94^. Of 4,055 micrographs, 3,099 micrographs were selected and 516 particles were manually picked. After 2D classification, 2D class averages were subjected to template-based auto-picking. 1,845,172 initial particles were picked and 4x binned. Multiple rounds of 2D classification left 927,902 particles, which produce 2D class averages showing secondary structure. An initial model was reconstructed with the selected particles in C1 symmetry and 3D classification further classified 330,509 particles. The selected 330,509 particles were re-extracted and unbinned to the original pixel (1.144 Å/pix). Initial 3D refinement reconstructed the cryo-EM map of DM3^Col-0^ at 4.53 Å resolution. CTF refinement and Bayesian polishing ^95^ improved the resolution to 3.57 Å. The polished particles were re-extracted again from micrographs containing 1 – 20 e^-^/Å^2^ dose. After multiple rounds of CTF refinement and Bayesian polishing, the particles reconstructed the cryo-EM map of DM3^Col-0^ at 3.39 Å resolution. All cryo-EM data processings were done using the software packages in SBgrid (https://sbgrid.org).

For cryo-EM data collection, 3 µL of the purified _ΔTP_DM3^Hh-0^ protein (0.2 mg/mL) was vitrified on the Quantifoil grid covered by GO. Cryo-EM data of _ΔTP_DM3^Hh-0^ were collected by a Titan Krios G4 (Thermo Fisher Scientific) with correlated double sampling (CDS) mode of K3 BioQuantum detector (Gatan), installed at Institute of Basic Science (IBS), Korea. Each movie frame contained a total dose of 49.8 e^-^/Å^2^ over 48 frames. 14,531 movie frames were obtained and motion correction was performed by Relion 3.1^96^. CTF estimation was accomplished by CTFFIND4^94^. 11,872 micrographs were selected and 532 particles were manually picked for the 2D template. The template-based autopicking picked 5,531,548 particles which were extracted and 4x binned. 2,982,001 particles were left after 2D classification and an initial model reconstructed in C1 symmetry. After 3D classification, 597,227 selected particles were re-extracted and unbinned to 0.849 Å/pix. Second 3D classification selected 413,056 particles and subsequent 3D refinement reconstructed a 3.53 Å resolution map. Multiple rounds of CTF refinement and Bayesian polishing improved the cryo-EM map of _ΔTP_DM3^Hh-0^ to 2.89 Å resolution^95^. To resolve the density of disrupted the dimer interface of DM3^Hh-0^, an optimal B-factor is determined by running MRC to MTZ from CCP-EM suite^94^ and applied to the cryo-EM map of _ΔTP_DM3^Hh-0^.

A detailed workflow and validation of cryo-EM maps of _ΔTP_DM3^Col-0^ (EMD-36240) and _ΔTP_DM3^Hh-0^ (EMD-36299) are illustrated in Figures S2 and S4. The distribution of local resolution was estimated by RESMAP implemented in Relion^97^.

### Structure determination and model validation

To build an atomic model of _ΔTP_DM3^Col-0^ (PDB ID: 8JGN), SWISS-MODEL^98^ generated an initial atomic model. The initial model was fitted into the 3.39 Å resolution cryo-EM map of _ΔTP_DM3^Col-0^ by using Chimera^98^. Real space refinement was performed by PHENIX^100^ and ISOLDE^101^. The model was further refined manually with Coot^102,103^. The real space refinement and manual refinement were repeated to improve model statistics, assessed by MolProbity^104^.

To determine the cryo-EM structure of _ΔTP_DM3^Hh-0^ (PDB ID: 8JGM), the cryo-EM structure of _ΔTP_DM3^Col-0^ was used for modelling with cryo-EM map of _ΔTP_DM3^Hh-0^. In the structure, T165 and T359 were substituted by I165 and A359, respectively. A part of the dimerizing loop (335-340) was not modelled due to the poor density (Figure S4). Real space refinement and manual refinement were repeated and performed by ISOLDE and Coot, respectively^101,102^. The model statistics of DM3^Hh-0^ was validated by MolProbity^104^. ChimeraX and PyMOL were used for model depiction and structure display^105^.

### Thermal shift assay

To measure thermal stability of the recombinant DM3 protein, 22.5 μl of 0.1mg/mL DM3 was mixed with 2.5 μL 10X SYPRO Orange dye (Thermo Fisher Scientific) to make 25 μL of reaction mixture. With an increment of 0.5 °C every 30 seconds, fluorescence signal was monitored by FRET mode of CFX Opus 96 Real-Time PCR System (Biorad) from 25 °C to 95 °C. Melting temperature (T_m_) was determined as the first derivative of RFU over temperature (-dF/dT) reaches the minimum value. T_m_ values of DM3 and the structure-guided mutants were calculated from an average T_m_ value from three individual experiments (n=3).

### Prolyl aminopeptidase activity assay

To measure prolyl aminopeptidase (PAP) activity of DM3, L-proline-7-amido-4-methylcoumarin hydrobromide (Pro-AMC, Merck) was used as an enzyme substrate. Pro-AMC was dissolved in distilled water to make 20µM stock concentration. 95 µL of 0.5 nM purified DM3 protein was prepared in a 96-well plate and 5 µL of 20 µM Pro-AMC was added to the plate before measurement. An increase of fluorescent signal from free AMC (λex 365nm; λem 440 nm) was monitored by using BioTek Synergy H1 (Agilent) at room temperature. For 1 hour, the fluorescent signal was measured every minute.

### Constructs for protein expression in *Nb*

In general, PCR amplification of sequences for cloning was performed using KOD-plus-Neo DNA polymerase (TOYOBO Ltd., Osaka, Japan). All oligonucleotides were supplied by Integrated DNA Technologies, Inc. (Coralville, IA) and are listed in Table S2. All restriction enzymes were supplied by Thermo Fisher Scientific. Following electrophoresis, PCR amplicons were purified using EZ-10 Spin Column Gel Extraction Kits (Bio Basic Asia Pacific Pte Ltd., Singapore). pJLSmart was used as the entry vector, while pGWB515, pGWB514, pGW512 were used as the destination vectors for N-terminus HA-tagged, C-terminus HA-tagged, and N-terminus FLAG-tagged DM3 constructs, respectively. Binary vector cloning was performed with Gateway™ LR Clonase™ II Enzyme mix (Thermo Fisher Scientific, Madison, WI). Site-directed mutagenesis was applied to generate mutations in DM3. All constructs were verified by Sanger sequencing. DM2h^Bla-1^ with an endogenous promoter was cloned into pMLBart for protein expression in plants.

### Plant materials and transient expression in *Nb*

*Nb* plants were grown at 24°C in a growth room under a 14 h light/10 h dark cycle. Five-week-old plants were used for *Agrobacterium tumefaciens*-mediated transient expression. *A. tumefaciens* (strain GV3101) carrying a binary construct was grown overnight at 28°C in an orbital shaker at 200 rpm/min in LB medium containing appropriate antibiotics to OD_600_ over 1. Bacterial cells were harvested and resuspended in the induction medium containing 10 mM MES pH 5.6, 10 mM MgCl_2_ and 500 μM acetosyringone with adjusted OD_600_ of 1. For HR assay, OD_600_ of 0.5 was used for DM3 and GFP; OD_600_ of 0.2 was used for DM2h. After 3 hours of incubation in induction media in the dark at room temperature, two bacterial inocula were mixed at a 1:1 volume ratio with the addition of 1/10 volume of an *Agrobacterium* culture carrying P19 to suppress gene silencing. Bacterial mixtures were manually infiltrated using a 1 mL needleless syringe at the abaxial side of the leaves. For DM3 protein detection with western blotting, OD_600_ of 0.5 was used for all constructs. Infiltrated leaves were kept in the dark wrapped with aluminium foils for two days, and photos of HR were taken around five-day post-infiltration.

### Plant protein extraction and western blot

150 mg leaf tissue was ground into fine powder in liquid nitrogen and homogenized in 200 μL of lysis buffer (0.33 M sucrose, 20 mM Tris-HCl pH 7.5, 1 mM EDTA, 10 mM DTT, and 1 x cOmplete™, EDTA-free Protease Inhibitor Cocktail (Roche, Madison, WI)). The lysate was cleared by a table-top centrifugation at 10,000 x g for 10 minutes at 4°C. The supernatant was mixed with 4x Laemmli sample buffer (240 mM Tris-HCl pH 6.8, 8% SDS, 40% glycerol, 5% β−mercaptoethanol, 0.04% Bromophenol blue), boiled for 10 minutes at 95°C, separated on 10% SDS-PAGE gel and transferred to PVDF membrane (Bio-Rad, Hercules, California). The membrane was incubated with anti-HA-Peroxidase (Sigma-Aldrich, 12013819001, clone 3F10) (1:2,500 dilution) overnight, followed by 3 times of TBS-T washing, for 15 minutes each time. Clarity Max Western ECL Substrate (Bio-Rad, Hercules, California) was used for chemiluminescence detection by Bio-Rad Chemidoc system.

### *In vivo* co-immunoprecipitation (co-IP) assay and blue native PAGE (BN-PAGE)

Protocol was modified from Ahn et al. ^106^. 0.5 g *Nb* leaf tissue was ground into fine powder in liquid nitrogen and homogenized in 1 mL of co-IP buffer (50 mM Tris-HCl, pH7.5, 50mM NaCl, 10% glycerol, 0.2% NP-40, 10 mM DTT and protease inhibitor cocktail (1 tablet/50mL)) by vortexing. Protein extract was incubated with anti-FLAG Affinity gel (Sigma-Aldrich)/ Myc-Trap Agarose (ChromoTek) for 1 hour at 4°C with rotation. The agarose was collected and washed 3 times with co-IP buffer. IP product was eluted by 100 uL of 150 nM FLAG pepdide or 0.1 μg/mL 2x Myc peptide with 1 hour incubation at 4°C. 10% of input is shown for FLAG-DM3 and DM2h^Bla-1^-Myc; 20% for HA-DM3. For BN-PAGE, IP products were diluted as per the manufacturer’s instructions by adding NativePAGE 5% G-250 sample additive, 4x Sample Buffer and water. Same amount of sample was run in SDS-PAGE as previously described as loading control. Samples were loaded and run on Native PAGE 4%-16% Bis-Tris gels alongside NativeMark unstained protein standard (Invitrogen). NativeMark unstained protein standard lane was cut for Coomassie blue staining. The proteins were then transferred to PVDF membranes. Proteins were fixed to the membranes by incubating with 8% acetic acid for 15 minutes, washed with water and left to dry. Membranes were subsequently re-activated with ethanol. Membranes were incubated in 1x SDS running buffer with 10 mM DTT for 1 hour, followed by 3 times 15 minutes TBS-T washing. The immunoprecipitated proteins and input proteins were analysed by immunoblotting with indicated antibodies as described. DM3 antibody from mouse was generated by Genscript using peptide CAKKEDWPPLYDVPR, 0.2 μg/mL of DM3 antibody was used for western blot.

### Purification of DM3 from *Nb*

*Nb-eds1* was used for infiltration to prevent cell death triggered by DM3^Hh-0^ and DM2h^Bla1^. Agrobacterium with desired constructs were 1:1 mixed and infiltrated to *Nb-eds1.* Approximately 20 g of samples for FLAG-_ΔTP_DM3^Hh-0^ and 35 g for FLAG-_ΔTP_DM3^Col-0^ were collected 40 hours after infiltration and ground into fine powder in liquid nitrogen. Plant extracts were resuspended using the co-IP buffer as described. The resuspended extracts were then sonicated using a hand-held sonicator (Sonics & Materials) with 2-second intervals, repeated 20 times. After centrifugation at 15,000 rpm for 30 minutes at 4°C, followed by 20 minutes using a high-speed vacuum centrifuge (Beckman Coulter), filtration was applied to get rid of plant debris. The filtered sample was incubated with anti-FLAG resin (GenScript) for 2 hours at 4°C on a rotator and the resins were washed with the resuspension buffer. FLAG-_ΔTP_DM3 was eluted using a resuspension buffer supplemented with 0.4 mg/mL FLAG peptide. Elution sample was concentrated at 3,300 rpm for 5 minutes per round until it reached 500 uL with Amicon Ultra-4 30K centrifugal filter system (Merck Millipore). Size exclusion chromatography was performed with Superdex200 increase 10/300 column (Cytiva) equilibrated with a gel filtration buffer (50 mM Tris-HCl pH 8.0, 100 mM NaCl). Elution fractions were concentrated using an Amicon Ultra-0.5 10K centrifugal filter system (Merck Millipore) and analysed by SDS-PAGE and 7% Tris-Glycine Native-PAGE (Separation gel: 4.6 mL 30% w/v acrylamide, 5 mL 1.5M Tris pH 8.8, 10.4 mL dH_2_O, APS 100 uL, TEMED 20 μL; stacking gel: 0.8 mL 30% w/v acrylamide, 2 mL 0.5M Tris pH 6.8, 5.2 mL dH_2_O, APS 60 uL, TEMED 10 μL; 5X sample buffer: 60 mM Tris-HCl pH6.8, 25% glycerol, 0.1% bromophenol blue) and western blot.

### Microscopic analyses: bimolecular fluorescence complementation

Vectors for BiFC originated from Kim et al.^107^ which are deposit in Addgene, plasmid number 48255, 48256, 48257, 48258. *Nb* leaves infiltrated with Agrobacteria mixture of 1:1 nYFP and cYFP were observed 30 hours after infiltration using Olympus FV3000 system and the images were processed using Olympus FV31S-SW. YFP were excited at 514 nm and obtained in 530-580 nm.

## QUANTIFICATION AND STATISTICAL ANALYSIS

Data for quantification analyses are presented as mean ± standard error (SE) or standard deviation (SD). The statistical analyses were performed by Student’s t-test or one-way analysis of variance (ANOVA) test (* P < 0.05, ** P < 0.01, *** P < 0.001, **** P < 0.0001, n.s., no significance). The number of replicates is shown in the figure legends.

## REFERENCES

1. Alcázar, R., Garcia, A.V., Parker, J.E., and Reymond, M. (2009). Incremental steps toward incompatibility revealed by Arabidopsis epistatic interactions modulating salicylic acid pathway activation. Proc Natl Acad Sci U S A 106, 334–339. 10.1073/pnas.0811734106.

2. Alcázar, R., von Reth, M., Bautor, J., Chae, E., Weigel, D., Koornneef, M., and Parker, J.E. (2014). Analysis of a plant complex resistance gene locus underlying immune-related hybrid incompatibility and its occurrence in nature. PLoS Genet 10, e1004848. 10.1371/journal.pgen.1004848.

3. Bomblies, K., Lempe, J., Epple, P., Warthmann, N., Lanz, C., Dangl, J.L., and Weigel, D. (2007). Autoimmune response as a mechanism for a Dobzhansky-Muller-type incompatibility syndrome in plants. PLoS biology 5, e236–e236. 10.1371/journal.pbio.0050236.

4. Chae, E., Bomblies, K., Kim, S.T., Karelina, D., Zaidem, M., Ossowski, S., Martin-Pizarro, C., Laitinen, R.A., Rowan, B.A., Tenenboim, H., et al. (2014). Species-wide genetic incompatibility analysis identifies immune genes as hot spots of deleterious epistasis. Cell 159, 1341–1351. 10.1016/j.cell.2014.10.049.

5. Ordon, J., Martin, P., Erickson, J.L., Ferik, F., Balcke, G., Bonas, U., and Stuttmann, J. (2021). Disentangling cause and consequence: genetic dissection of the DANGEROUS MIX2 risk locus, and activation of the DM2h NLR in autoimmunity. Plant J 106, 1008–1023. 10.1111/tpj.15215.

6. Atanasov, K.E., Liu, C., Erban, A., Kopka, J., Parker, J.E., and Alcazar, R. (2018). NLR Mutations Suppressing Immune Hybrid Incompatibility and Their Effects on Disease Resistance. Plant Physiol 177, 1152–1169. 10.1104/pp.18.00462.

7. Jones, J.D.G., Vance, R.E., and Dangl, J.L. (2016). Intracellular innate immune surveillance devices in plants and animals. Science 354, aaf6395. doi:10.1126/science.aaf6395.

8. Kourelis, J., and van der Hoorn, R.A.L. (2018). Defended to the Nines: 25 Years of Resistance Gene Cloning Identifies Nine Mechanisms for R Protein Function. Plant Cell 30, 285–299. 10.1105/tpc.17.00579.

9. Kourelis, J., Sakai, T., Adachi, H., and Kamoun, S. (2021). RefPlantNLR is a comprehensive collection of experimentally validated plant disease resistance proteins from the NLR family. PLoS Biol 19, e3001124. 10.1371/journal.pbio.3001124.

10. Ao, K., and Li, X. (2022). Indirect recognition of pathogen effectors by NLRs. Essays Biochem 66, 485–500. 10.1042/ebc20210097.

11. Dangl, J.L., and Jones, J.D. (2001). Plant pathogens and integrated defence responses to infection. Nature 411, 826–833. 10.1038/35081161.

12. Van Der Biezen, E.A., and Jones, J.D.G. (1998). Plant disease-resistance proteins and the gene-for-gene concept. Trends in Biochem Sciences 23, 454–456. 10.1016/S0968-0004(98)01311-5.

13. van der Hoorn, R.A., and Kamoun, S. (2008). From Guard to Decoy: a new model for perception of plant pathogen effectors. Plant Cell 20, 2009–2017. 10.1105/tpc.108.060194.

14. Ade, J., DeYoung, B.J., Golstein, C., and Innes, R.W. (2007). Indirect activation of a plant nucleotide binding site-leucine-rich repeat protein by a bacterial protease. Proc Natl Acad Sci U S A 104, 2531–2536. 10.1073/pnas.0608779104.

15. Chung, E.H., da Cunha, L., Wu, A.J., Gao, Z., Cherkis, K., Afzal, A.J., Mackey, D., and Dangl, J.L. (2011). Specific threonine phosphorylation of a host target by two unrelated type III effectors activates a host innate immune receptor in plants. Cell Host Microbe 9, 125–136. 10.1016/j.chom.2011.01.009.

16. Kim, H., Ahn, Y.J., Lee, H., Chung, E.H., Segonzac, C., and Sohn, K.H. (2023). Diversified host target families mediate convergently evolved effector recognition across plant species. Curr Opin Plant Biol 74, 102398. 10.1016/j.pbi.2023.102398.

17. Kim, M.G., da Cunha, L., McFall, A.J., Belkhadir, Y., DebRoy, S., Dangl, J.L., and Mackey, D. (2005). Two Pseudomonas syringae type III effectors inhibit RIN4-regulated basal defense in Arabidopsis. Cell 121, 749–759. 10.1016/j.cell.2005.03.025.

18. Mackey, D., Belkhadir, Y., Alonso, J.M., Ecker, J.R., and Dangl, J.L. (2003). Arabidopsis RIN4 is a target of the type III virulence effector AvrRpt2 and modulates RPS2-mediated resistance. Cell 112, 379–389. 10.1016/s0092-8674(03)00040-0.

19. Mackey, D., Holt, B.F., 3rd, Wiig, A., and Dangl, J.L. (2002). RIN4 interacts with Pseudomonas syringae type III effector molecules and is required for RPM1-mediated resistance in Arabidopsis. Cell 108, 743–754. 10.1016/s0092-8674(02)00661-x.

20. Qi, D., DeYoung, B.J., and Innes, R.W. (2012). Structure-function analysis of the coiled-coil and leucine-rich repeat domains of the RPS5 disease resistance protein. Plant Physiol 158, 1819–1832. 10.1104/pp.112.194035.

21. Ma, S., Lapin, D., Liu, L., Sun, Y., Song, W., Zhang, X., Logemann, E., Yu, D., Wang, J., Jirschitzka, J., et al. (2020). Direct pathogen-induced assembly of an NLR immune receptor complex to form a holoenzyme. Science 370, eabe3069. 10.1126/science.abe3069.

22. Martin, R., Qi, T., Zhang, H., Liu, F., King, M., Toth, C., Nogales, E., and Staskawicz, B.J. (2020). Structure of the activated ROQ1 resistosome directly recognizing the pathogen effector XopQ. Science 370, eabd9993. 10.1126/science.abd9993.

23. Wang, J., Wang, J., Hu, M., Wu, S., Qi, J., Wang, G., Han, Z., Qi, Y., Gao, N., Wang, H.-W., et al. (2019). Ligand-triggered allosteric ADP release primes a plant NLR complex. Science 364, eaav5868. 10.1126/science.aav5868.

24. Wang, J., Hu, M., Wang, J., Qi, J., Han, Z., Wang, G., Qi, Y., Wang, H.-W., Zhou, J.-M., and Chai, J. (2019). Reconstitution and structure of a plant NLR resistosome conferring immunity. Science 364, eaav5870. 10.1126/science.aav5870.

25. Gong, Y., Tian, L., Kontos, I., Li, J., and Li, X. (2023). Plant immune signaling network mediated by helper NLRs. Curr Opin Plant Biol 73, 102354. 10.1016/j.pbi.2023.102354.

26. Jia, A., Huang, S., Song, W., Wang, J., Meng, Y., Sun, Y., Xu, L., Laessle, H., Jirschitzka, J., Hou, J., et al. (2022). TIR-catalyzed ADP-ribosylation reactions produce signaling molecules for plant immunity. Science 377, eabq8180. 10.1126/science.abq8180.

27. Yu, D., Song, W., Tan, E.Y.J., Liu, L., Cao, Y., Jirschitzka, J., Li, E., Logemann, E., Xu, C., Huang, S., et al. (2022). TIR domains of plant immune receptors are 2’,3’-cAMP/cGMP synthetases mediating cell death. Cell 185, 2370–2386.e2318. 10.1016/j.cell.2022.04.032.

28. Bayless, A.M., Chen, S., Ogden, S.C., Xu, X., Sidda, J.D., Manik, M.K., Li, S., Kobe, B., Ve, T., Song, L., et al. (2023). Plant and prokaryotic TIR domains generate distinct cyclic ADPR NADase products. Sci Adv 9, eade8487. 10.1126/sciadv.ade8487.

29. Horsefield, S., Burdett, H., Zhang, X., Manik, M.K., Shi, Y., Chen, J., Qi, T., Gilley, J., Lai, J.S., Rank, M.X., et al. (2019). NAD(+) cleavage activity by animal and plant TIR domains in cell death pathways. Science 365, 793–799. 10.1126/science.aax1911.

30. Manik, M.K., Shi, Y., Li, S., Zaydman, M.A., Damaraju, N., Eastman, S., Smith, T.G., Gu, W., Masic, V., Mosaiab, T., et al. (2022). Cyclic ADP ribose isomers: Production, chemical structures, and immune signaling. Science 377, eadc8969. 10.1126/science.adc8969.

31. Wan, L., Essuman, K., Anderson, R.G., Sasaki, Y., Monteiro, F., Chung, E.H., Osborne Nishimura, E., DiAntonio, A., Milbrandt, J., Dangl, J.L., and Nishimura, M.T. (2019). TIR domains of plant immune receptors are NAD(+)-cleaving enzymes that promote cell death. Science 365, 799–803. 10.1126/science.aax1771.

32. Huang, S., Jia, A., Song, W., Hessler, G., Meng, Y., Sun, Y., Xu, L., Laessle, H., Jirschitzka, J., Ma, S., et al. (2022). Identification and receptor mechanism of TIR-catalyzed small molecules in plant immunity. Science 377, eabq3297. 10.1126/science.abq3297.

33. van Wersch, R., Li, X., and Zhang, Y. (2016). Mighty Dwarfs: Arabidopsis Autoimmune Mutants and Their Usages in Genetic Dissection of Plant Immunity. Front Plant Sci 7. 10.3389/fpls.2016.01717.

34. Freh, M., Gao, J., Petersen, M., and Panstruga, R. (2022). Plant autoimmunity-fresh insights into an old phenomenon. Plant Physiol 188, 1419–1434. 10.1093/plphys/kiab590.

35. Kourelis, J., and Adachi, H. (2022). Activation and regulation of NLR immune receptor networks. Plant Cell Physiol 63, 1366–1377. 10.1093/pcp/pcac116.

36. Wan, W.L., Kim, S.T., Castel, B., Charoennit, N., and Chae, E. (2021). Genetics of autoimmunity in plants: an evolutionary genetics perspective. New Phytol 229, 1215–1233. 10.1111/nph.16947.

37. Alcázar, R., García, A.V., Kronholm, I., de Meaux, J., Koornneef, M., Parker, J.E., and Reymond, M. (2010). Natural variation at Strubbelig Receptor Kinase 3 drives immune-triggered incompatibilities between Arabidopsis thaliana accessions. Nat Genet 42, 1135–1139. 10.1038/ng.704.

38. Platre, M.P., Satbhai, S.B., Brent, L., Gleason, M.F., Cao, M., Grison, M., Glavier, M., Zhang, L., Gaillochet, C., Goeschl, C., et al. (2022). The receptor kinase SRF3 coordinates iron-level and flagellin dependent defense and growth responses in plants. Nat Commun 13, 4445. 10.1038/s41467-022-32167-6.

39. Tahir, J., Watanabe, M., Jing, H.-C., Hunter, D.A., Tohge, T., Nunes-Nesi, A., Brotman, Y., Fernie, A.R., Hoefgen, R., and Dijkwel, P.P. (2013). Activation of R-mediated innate immunity and disease susceptibility is affected by mutations in a cytosolic O-acetylserine (thiol) lyase in Arabidopsis. The Plant Journal 73, 118–130. 10.1111/tpj.12021.

40. Shirzadian-Khorramabad, R., Jing, H.C., Everts, G.E., Schippers, J.H., Hille, J., and Dijkwel, P.P. (2010). A mutation in the cytosolic O-acetylserine (thiol) lyase induces a genome-dependent early leaf death phenotype in Arabidopsis. BMC Plant Biol 10, 80. 10.1186/1471-2229-10-80.

41. Stuttmann, J., Peine, N., Garcia, A.V., Wagner, C., Choudhury, S.R., Wang, Y., James, G.V., Griebel, T., Alcázar, R., Tsuda, K., et al. (2016). Arabidopsis thaliana DM2h (R8) within the Landsberg RPP1-like resistance locus underlies three different cases of EDS1-conditioned autoimmunity. PLoS Genet 12, e1005990. 10.1371/journal.pgen.1005990.

42. Ghifari, A.S., Teixeira, P.F., Kmiec, B., Pružinská, A., Glaser, E., and Murcha, M.W. (2020). A mitochondrial prolyl aminopeptidase PAP2 releases N-terminal proline and regulates proline homeostasis during stress response. Plant J 104, 1182–1194. 10.1111/tpj.14987.

43. Ghifari, A.S., Teixeira, P.F., Kmiec, B., Singh, N., Glaser, E., and Murcha, M.W. (2022). The dual-targeted prolyl aminopeptidase PAP1 is involved in proline accumulation in response to stress and during pollen development. Journal of Experimental Botany 73, 78–93. 10.1093/jxb/erab397.

44. The 1001 Genomes Consortium. (2016). 1,135 Genomes Reveal the Global Pattern of Polymorphism in Arabidopsis thaliana. Cell 166, 481–491. 10.1016/j.cell.2016.05.063.

45. Bauer, T.L., Buchholz, P.C.F., and Pleiss, J. (2020). The modular structure of alpha/beta-hydrolases. FEBS J 287, 1035–1053. 10.1111/febs.15071.

46. Mindrebo, J.T., Nartey, C.M., Seto, Y., Burkart, M.D., and Noel, J.P. (2016). Unveiling the functional diversity of the alpha/beta hydrolase superfamily in the plant kingdom. Curr Opin Struct Biol 41, 233–246. 10.1016/j.sbi.2016.08.005.

47. Zhang, X., Bernoux, M., Bentham, A.R., Newman, T.E., Ve, T., Casey, L.W., Raaymakers, T.M., Hu, J., Croll, T.I., Schreiber, K.J., et al. (2017). Multiple functional self-association interfaces in plant TIR domains. Proc Natl Acad Sci U S A 114, E2046–e2052. 10.1073/pnas.1621248114.

48. Williams, S.J., Sohn, K.H., Wan, L., Bernoux, M., Sarris, P.F., Segonzac, C., Ve, T., Ma, Y., Saucet, S.B., Ericsson, D.J., et al. (2014). Structural Basis for Assembly and Function of a Heterodimeric Plant Immune Receptor. Science 344, 299–303. doi:10.1126/science.1247357.

49. Jiao, W.B., and Schneeberger, K. (2020). Chromosome-level assemblies of multiple Arabidopsis genomes reveal hotspots of rearrangements with altered evolutionary dynamics. Nat Commun 11, 989. 10.1038/s41467-020-14779-y.

50. Lee, R.R.Q., and Chae, E. (2020). Variation patterns of NLR clusters in Arabidopsis thaliana genomes. Plant Commun 1, 100089. 10.1016/j.xplc.2020.100089.

51. Prigozhin, D.M., and Krasileva, K.V. (2021). Analysis of intraspecies diversity reveals a subset of highly variable plant immune receptors and predicts their binding sites. Plant Cell 33, 998–1015. 10.1093/plcell/koab013.

52. Botos, I., Lountos, G.T., Wu, W., Cherry, S., Ghirlando, R., Kudzhaev, A.M., Rotanova, T.V., de Val, N., Tropea, J.E., Gustchina, A., and Wlodawer, A. (2019). Cryo-EM structure of substrate-free E. coli Lon protease provides insights into the dynamics of Lon machinery. Curr Res Struct Biol 1, 13–20. 10.1016/j.crstbi.2019.10.001.

53. Brandstetter, H., Kim, J.S., Groll, M., and Huber, R. (2001). Crystal structure of the tricorn protease reveals a protein disassembly line. Nature 414, 466–470. 10.1038/35106609.

54. García-Nafría, J., Ondrovicová, G., Blagova, E., Levdikov, V.M., Bauer, J.A., Suzuki, C.K., Kutejová, E., Wilkinson, A.J., and Wilson, K.S. (2010). Structure of the catalytic domain of the human mitochondrial Lon protease: proposed relation of oligomer formation and activity. Protein Sci 19, 987–999. 10.1002/pro.376.

55. Kirthika, P., Lloren, K.K.S., Jawalagatti, V., and Lee, J.H. (2023). Structure, Substrate Specificity and Role of Lon Protease in Bacterial Pathogenesis and Survival. Int J Mol Sci 24. 10.3390/ijms24043422.

56. Shin, M., Puchades, C., Asmita, A., Puri, N., Adjei, E., Wiseman, R.L., Karzai, A.W., and Lander, G.C. (2020). Structural basis for distinct operational modes and protease activation in AAA+ protease Lon. Sci Adv 6, eaba8404. 10.1126/sciadv.aba8404.

57. Tamura, T., Tamura, N., Cejka, Z., Hegerl, R., Lottspeich, F., and Baumeister, W. (1996). Tricorn protease--the core of a modular proteolytic system. Science 274, 1385–1389. 10.1126/science.274.5291.1385.

58. Kim, G., Azmi, L., Jang, S., Jung, T., Hebert, H., Roe, A.J., Byron, O., and Song, J.-J. (2019). Aldehyde-alcohol dehydrogenase forms a high-order spirosome architecture critical for its activity. Nat Commun 10, 4527. 10.1038/s41467-019-12427-8.

59. Lynch, E.M., and Kollman, J.M. (2020). Coupled structural transitions enable highly cooperative regulation of human CTPS2 filaments. Nat. Struct. Mol. Biol. 27, 42–48. 10.1038/s41594-019-0352-5.

60. Lynch, E.M., Kollman, J.M., and Webb, B.A. (2020). Filament formation by metabolic enzymes-A new twist on regulation. Curr Opin Cell Biol 66, 28–33. 10.1016/j.ceb.2020.04.006.

61. Hvorecny, K.L., and Kollman, J.M. (2023). Greater than the sum of parts: Mechanisms of metabolic regulation by enzyme filaments. Curr Opin Struct Biol 79, 102530. 10.1016/j.sbi.2023.102530.

62. Bhandari, D.D., Lapin, D., Kracher, B., von Born, P., Bautor, J., Niefind, K., and Parker, J.E. (2019). An EDS1 heterodimer signalling surface enforces timely reprogramming of immunity genes in Arabidopsis. Nat Commun 10, 772. 10.1038/s41467-019-08783-0.

63. Li, L., and Weigel, D. (2021). One Hundred Years of Hybrid Necrosis: Hybrid Autoimmunity as a Window into the Mechanisms and Evolution of Plant–Pathogen Interactions. Annu Rev Phytopathol 59, 213–237. 10.1146/annurev-phyto-020620-114826.

64. Tran, D.T.N., Chung, E.H., Habring-Muller, A., Demar, M., Schwab, R., Dangl, J.L., Weigel, D., and Chae, E. (2017). Activation of a Plant NLR Complex through Heteromeric Association with an Autoimmune Risk Variant of Another NLR. Curr Biol 27, 1148–1160. 10.1016/j.cub.2017.03.018.

65. Alvarez, C., Bermudez, M.A., Romero, L.C., Gotor, C., and Garcia, I. (2012). Cysteine homeostasis plays an essential role in plant immunity. New Phytol 193, 165–177. 10.1111/j.1469-8137.2011.03889.x.

66. Alvarez, M.E., Savouré, A., and Szabados, L. (2022). Proline metabolism as regulatory hub. Trends Plant Sci 27, 39–55. 10.1016/j.tplants.2021.07.009.

67. Orth-He, E.L., Huang, H.C., Rao, S.D., Wang, Q., Chen, Q., O’Mara, C.M., Chui, A.J., Saoi, M., Griswold, A.R., Bhattacharjee, A., et al. (2023). Protein folding stress potentiates NLRP1 and CARD8 inflammasome activation. Cell Rep 42, 111965. 10.1016/j.celrep.2022.111965.

68. Rao, S.D., Chen, Q., Wang, Q., Orth-He, E.L., Saoi, M., Griswold, A.R., Bhattacharjee, A., Ball, D.P., Huang, H.-C., Chui, A.J., et al. (2022). M24B aminopeptidase inhibitors selectively activate the CARD8 inflammasome. Nat Chem Biol 18, 565–574. 10.1038/s41589-021-00964-7.

69. Li, L., Habring, A., Wang, K., and Weigel, D. (2020). Atypical Resistance Protein RPW8/HR Triggers Oligomerization of the NLR Immune Receptor RPP7 and Autoimmunity. Cell Host & Microbe 27, 405–417.e406. 10.1016/j.chom.2020.01.012.

70. Selvaraj, M., Toghani, A., Pai, H., Sugihara, Y., Kourelis, J., Yuen, E.L.H., Ibrahim, T., Zhao, H., Xie, R., Maqbool, A., et al. (2023). Activation of plant immunity through conversion of a helper NLR homodimer into a resistosome. bioRxiv, 2023.2012.2017.572070. 10.1101/2023.12.17.572070.

71. Schreiber, K.J., Bentham, A., Williams, S.J., Kobe, B., and Staskawicz, B.J. (2016). Multiple Domain Associations within the Arabidopsis Immune Receptor RPP1 Regulate the Activation of Programmed Cell Death. PLoS Pathog 12, e1005769. 10.1371/journal.ppat.1005769.

72. Steinbrenner, A.D., Goritschnig, S., and Staskawicz, B.J. (2015). Recognition and Activation Domains Contribute to Allele-Specific Responses of an Arabidopsis NLR Receptor to an Oomycete Effector Protein. PLoS Pathog 11, e1004665. 10.1371/journal.ppat.1004665.

73. McLellan, H., Harvey, S.E., Steinbrenner, J., Armstrong, M.R., He, Q., Clewes, R., Pritchard, L., Wang, W., Wang, S., Nussbaumer, T., et al. (2022). Exploiting breakdown in nonhost effector-target interactions to boost host disease resistance. Proc Natl Acad Sci U S A 119, e2114064119. 10.1073/pnas.2114064119.

74. Bastedo, D.P., Khan, M., Martel, A., Seto, D., Kireeva, I., Zhang, J., Masud, W., Millar, D., Lee, J.Y., Lee, A.H., et al. (2019). Perturbations of the ZED1 pseudokinase activate plant immunity. PLoS Pathog 15, e1007900. 10.1371/journal.ppat.1007900.

75. Choi, S., Prokchorchik, M., Lee, H., Gupta, R., Lee, Y., Chung, E.H., Cho, B., Kim, M.S., Kim, S.T., and Sohn, K.H. (2021). Direct acetylation of a conserved threonine of RIN4 by the bacterial effector HopZ5 or AvrBsT activates RPM1-dependent immunity in Arabidopsis. Mol Plant 14, 1951–1960. 10.1016/j.molp.2021.07.017.

76. Lee, J., Manning, A.J., Wolfgeher, D., Jelenska, J., Cavanaugh, K.A., Xu, H., Fernandez, S.M., Michelmore, R.W., Kron, S.J., and Greenberg, J.T. (2015). Acetylation of an NB-LRR Plant Immune-Effector Complex Suppresses Immunity. Cell Rep 13, 1670–1682. 10.1016/j.celrep.2015.10.029.

77. Ma, K.W., Jiang, S., Hawara, E., Lee, D., Pan, S., Coaker, G., Song, J., and Ma, W. (2015). Two serine residues in Pseudomonas syringae effector HopZ1a are required for acetyltransferase activity and association with the host co-factor. New Phytol 208, 1157–1168. 10.1111/nph.13528.

78. Jeleńska, J., Lee, J., Manning, A.J., Wolfgeher, D.J., Ahn, Y., Walters-Marrah, G., Lopez, I.E., Garcia, L., McClerklin, S.A., Michelmore, R.W., et al. (2021). Pseudomonas syringae effector HopZ3 suppresses the bacterial AvrPto1-tomato PTO immune complex via acetylation. PLoS Pathog 17, e1010017. 10.1371/journal.ppat.1010017.

79. Jiang, S., Yao, J., Ma, K.W., Zhou, H., Song, J., He, S.Y., and Ma, W. (2013). Bacterial effector activates jasmonate signaling by directly targeting JAZ transcriptional repressors. PLoS Pathog 9, e1003715. 10.1371/journal.ppat.1003715.

80. Lewis, J.D., Lee, A.H., Hassan, J.A., Wan, J., Hurley, B., Jhingree, J.R., Wang, P.W., Lo, T., Youn, J.Y., Guttman, D.S., and Desveaux, D. (2013). The Arabidopsis ZED1 pseudokinase is required for ZAR1-mediated immunity induced by the Pseudomonas syringae type III effector HopZ1a. Proc Natl Acad Sci U S A 110, 18722–18727. 10.1073/pnas.1315520110.

81. Zhang, Z.M., Ma, K.W., Gao, L., Hu, Z., Schwizer, S., Ma, W., and Song, J. (2017). Mechanism of host substrate acetylation by a YopJ family effector. Nat Plants 3, 17115. 10.1038/nplants.2017.115.

82. Choi, S., Jayaraman, J., and Sohn, K.H. (2018). Arabidopsis thaliana SOBER1 (SUPPRESSOR OF AVRBST-ELICITED RESISTANCE 1) suppresses plant immunity triggered by multiple bacterial acetyltransferase effectors. New Phytol 219, 324–335. 10.1111/nph.15125.

83. Remick, B.C., Gaidt, M.M., and Vance, R.E. (2023). Effector-Triggered Immunity. Annual Review of Immunology 41, 453–481. 10.1146/annurev-immunol-101721-031732.

84. Ordon, J., Gantner, J., Kemna, J., Schwalgun, L., Reschke, M., Streubel, J., Boch, J., and Stuttmann, J. (2017). Generation of chromosomal deletions in dicotyledonous plants employing a user-friendly genome editing toolkit. Plant J 89, 155–168. 10.1111/tpj.13319.

85. Sperschneider, J., Catanzariti, A.M., DeBoer, K., Petre, B., Gardiner, D.M., Singh, K.B., Dodds, P.N., and Taylor, J.M. (2017). LOCALIZER: subcellular localization prediction of both plant and effector proteins in the plant cell. Sci Rep 7, 44598. 10.1038/srep44598.

86. Kumar, S., Stecher, G., Li, M., Knyaz, C., and Tamura, K. (2018). MEGA X: Molecular Evolutionary Genetics Analysis across Computing Platforms. Mol Biol Evol 35, 1547–1549. 10.1093/molbev/msy096.

87. Thompson, J.D., Higgins, D.G., and Gibson, T.J. (1994). CLUSTAL W: improving the sensitivity of progressive multiple sequence alignment through sequence weighting, position-specific gap penalties and weight matrix choice. Nucleic Acids Res 22, 4673–4680. 10.1093/nar/22.22.4673.

88. Chang, J.M., Di Tommaso, P., Taly, J.F., and Notredame, C. (2012). Accurate multiple sequence alignment of transmembrane proteins with PSI-Coffee. BMC Bioinformatics 13 *Suppl 4*, S1. 10.1186/1471-2105-13-s4-s1.

89. Larkin, M.A., Blackshields, G., Brown, N.P., Chenna, R., McGettigan, P.A., McWilliam, H., Valentin, F., Wallace, I.M., Wilm, A., Lopez, R., et al. (2007). Clustal W and Clustal X version 2.0. Bioinformatics 23, 2947–2948. 10.1093/bioinformatics/btm404.

90. Team, R.C. (2021). R: A Language and Environment for Statistical Computing. https://www.R-project.org/.

91. Paradis, E. (2010). pegas: an R package for population genetics with an integrated-modular approach. Bioinformatics 26, 419–420. 10.1093/bioinformatics/btp696.

92. Patel, A., Toso, D., Litvak, A., and Nogales, E. (2021). Efficient graphene oxide coating improves cryo-EM sample preparation and data collection from tilted grids. bioRxiv, 2021.2003.2008.434344. 10.1101/2021.03.08.434344.

93. Zheng, S.Q., Palovcak, E., Armache, J.-P., Verba, K.A., Cheng, Y., and Agard, D.A. (2017). MotionCor2: anisotropic correction of beam-induced motion for improved cryo-electron microscopy. Nat Methods 14, 331–332. 10.1038/nmeth.4193.

94. Zhang, K. (2016). Gctf: Real-time CTF determination and correction. Journal of Structural Biology 193, 1–12. 10.1016/j.jsb.2015.11.003.

95. Zivanov, J., Nakane, T., and Scheres, S.H.W. (2019). A Bayesian approach to beam-induced motion correction in cryo-EM single-particle analysis. IUCrJ 6, 5–17. 10.1107/S205225251801463X.

96. Scheres, S.H. (2012). RELION: implementation of a Bayesian approach to cryo-EM structure determination. J Struct Biol 180, 519–530. 10.1016/j.jsb.2012.09.006.

97. Kucukelbir, A., Sigworth, F.J., and Tagare, H.D. (2014). Quantifying the local resolution of cryo-EM density maps. Nat Methods 11, 63–65. 10.1038/nmeth.2727.

98. Waterhouse, A., Bertoni, M., Bienert, S., Studer, G., Tauriello, G., Gumienny, R., Heer, F.T., de Beer, T.A.P., Rempfer, C., Bordoli, L., et al. (2018). SWISS-MODEL: homology modelling of protein structures and complexes. Nucleic Acids Res 46, W296–W303. 10.1093/nar/gky427.

99. Pettersen, E.F., Goddard, T.D., Huang, C.C., Couch, G.S., Greenblatt, D.M., Meng, E.C., and Ferrin, T.E. (2004). UCSF Chimera--a visualization system for exploratory research and analysis. J Comput Chem 25, 1605–1612. 10.1002/jcc.20084.

100. Adams, P.D., Afonine, P.V., Bunkóczi, G., Chen, V.B., Davis, I.W., Echols, N., Headd, J.J., Hung, L.W., Kapral, G.J., Grosse-Kunstleve, R.W., et al. (2010). PHENIX: a comprehensive Python-based system for macromolecular structure solution. Acta Crystallogr D Biol Crystallogr 66, 213–221. 10.1107/s0907444909052925.

101. Croll, T.I. (2018). ISOLDE: a physically realistic environment for model building into low-resolution electron-density maps. Acta Crystallogr D Struct Biol 74, 519–530. 10.1107/s2059798318002425.

102. Emsley, P., Lohkamp, B., Scott, W.G., and Cowtan, K. (2010). Features and development of Coot. Acta Crystallogr D Biol Crystallogr 66, 486–501. 10.1107/S0907444910007493.

103. Goddard, T.D., Huang, C.C., Meng, E.C., Pettersen, E.F., Couch, G.S., Morris, J.H., and Ferrin, T.E. (2018). UCSF ChimeraX: Meeting modern challenges in visualization and analysis. Protein Sci 27, 14–25. 10.1002/pro.3235.

104. Davis, I.W., Leaver-Fay, A., Chen, V.B., Block, J.N., Kapral, G.J., Wang, X., Murray, L.W., Arendall, W.B., 3rd, Snoeyink, J., Richardson, J.S., and Richardson, D.C. (2007). MolProbity: all-atom contacts and structure validation for proteins and nucleic acids. Nucleic Acids Res 35, W375–383. 10.1093/nar/gkm216.

105. The PyMOL Molecular Graphics System, Version 1.3, Schrödinger, LLC.

106. Ahn, H.K., Lin, X., Olave-Achury, A.C., Derevnina, L., Contreras, M.P., Kourelis, J., Wu, C.H., Kamoun, S., and Jones, J.D.G. (2023). Effector-dependent activation and oligomerization of plant NRC class helper NLRs by sensor NLR immune receptors Rpi-amr3 and Rpi-amr1. Embo j 42, e111484. 10.15252/embj.2022111484.

107. Kim, J.-G., Li, X., Roden, J.A., Taylor, K.W., Aakre, C.D., Su, B., Lalonde, S., Kirik, A., Chen, Y., Baranage, G., et al. (2009). Xanthomonas T3S Effector XopN Suppresses PAMP-Triggered Immunity and Interacts with a Tomato Atypical Receptor-Like Kinase and TFT1. The Plant Cell 21, 1305–1323. 10.1105/tpc.108.063123.

