## Supplementary material for "Structure determinants of DANGEROUS MIX 3, an alpha/beta hydrolase, for triggering NLR-mediated genetic incompatibility in plants": Figure S1

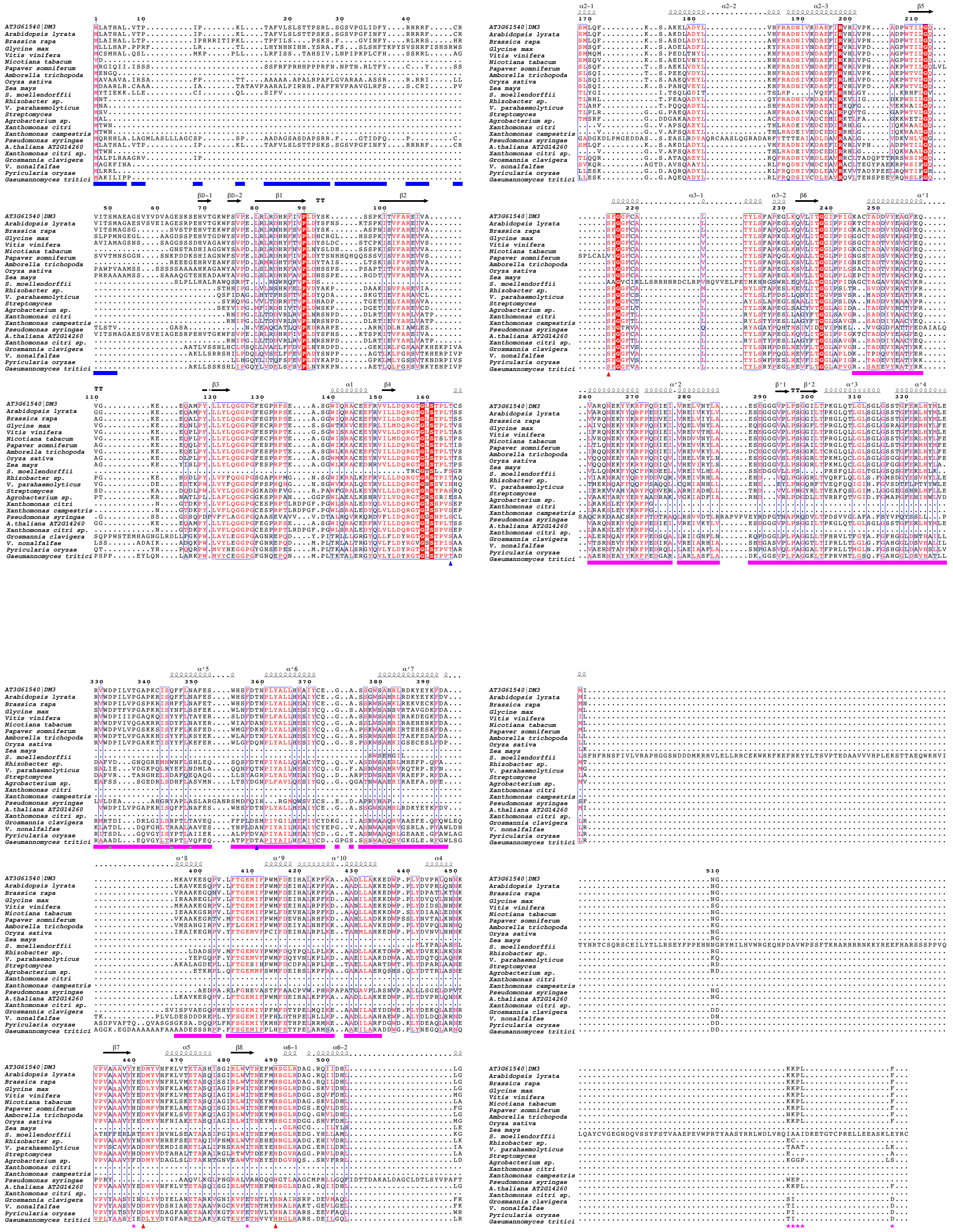

**Figure S1. Protein Sequence alignment and the secondary structure of DM3 homologs. Related to Figure 1.** The alignment portrays conservation between DM3$^{\text{Col-0}}$ homologs with indicated important residues. T165 and T359 are labeled with blue triangles to display important DM3 variant residues. Predicted catalytic triad residues S214, D462, and H490 are labeled with red triangles. Based on the DM3 structure in this study, the green star beneath indicates the important dimer interface residues: D332, Q345, and N349; the pink star indicates the trimer interface residues Y460, V484, and C-terminal K511-F515 of the DM3 hexameric complex. The blue box below indicates the N-terminal deletion of DM3$^{\text{Col-0}}$ (M1-M51) designed for structural studies. The magenta box below indicates the lid domain region. The protein sequence alignment was done using homology extension (PSI-Coffee) on the T-Coffee server.
