## Supplementary material for "Structure determinants of DANGEROUS MIX 3, an alpha/beta hydrolase, for triggering NLR-mediated genetic incompatibility in plants": Figure S2

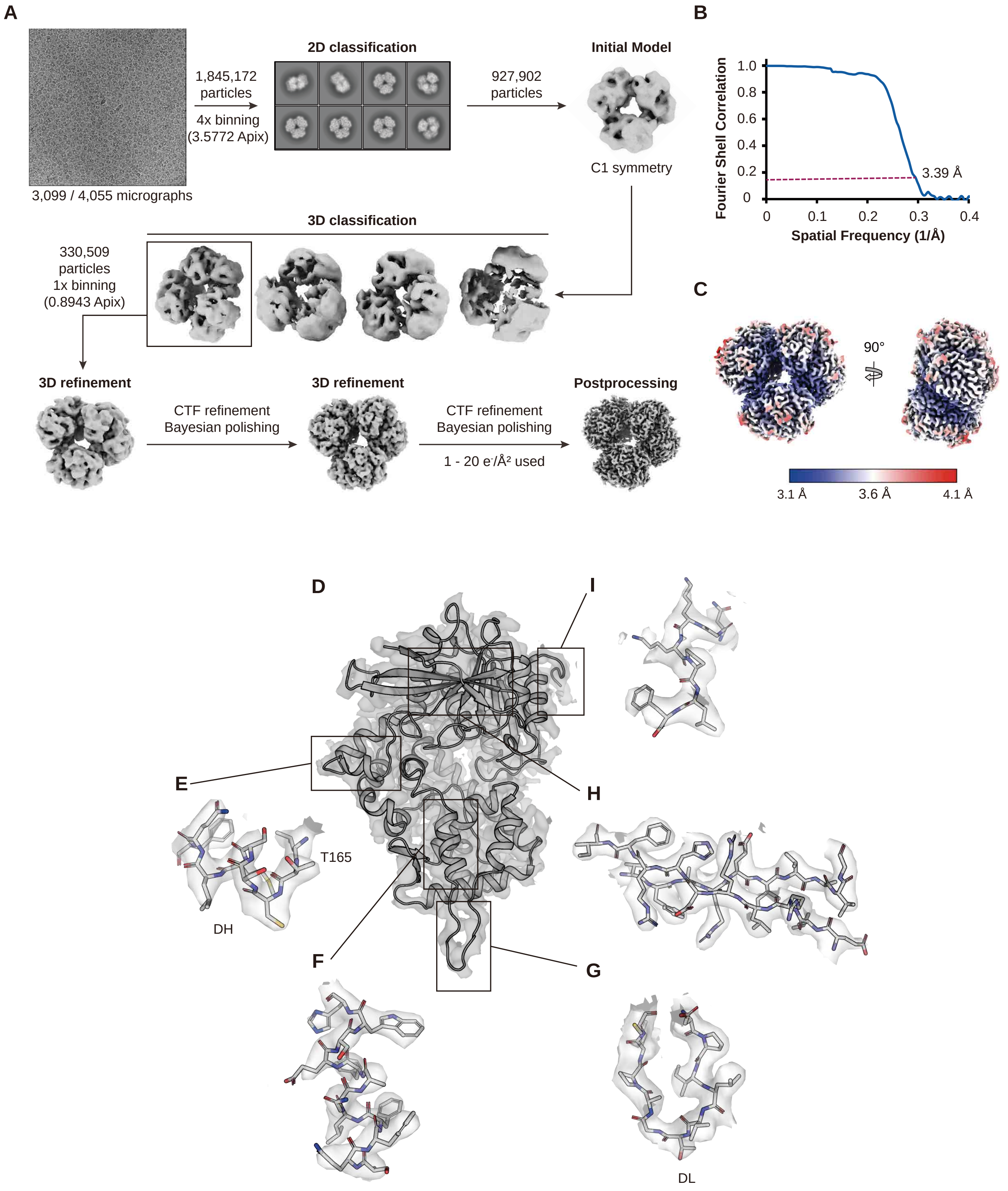


**Figure S2. Cryo-EM image processing on $_{\Delta TP}$DM3$^{Col\text{-}0}$ data and validation on EM map reconstruction. Related to Figure 2.** (A) A flowchart of the $_{\Delta TP}$DM3$^{Col\text{-}0}$ cryo-EM data processing is shown in detail. A representative cryo-EM micrograph, 2D class average, 3D classes and the final EM map (bottom right) are shown. Detailed descriptions of the data processing are in STAR Methods. (B) A FSC curve shows the resolution of the cryo-EM reconstruction of $_{\Delta TP}$DM3$^{Col\text{-}0}$ as 3.39 Å with 0.143 criteria. (C) Local resolution of the cryo-EM map of $_{\Delta TP}$DM3$^{Col\text{-}0}$ is estimated and colored by Relion. The scale bar is shown at the bottom. (D) The cryo-EM structure of $_{\Delta TP}$DM3$^{Col\text{-}0}$ monomer is fitted into the cryo-EM map. The structure of $_{\Delta TP}$DM3$^{Col\text{-}0}$ monomer and the density are colored dark grey and grey respectively. (E) A structure near the DH (164-172) is fitted in the cryo-EM. T165 is shown with the DH (166-171). (F) A α-helix of lid domain (344-355) is fitted in the cryo-EM density. (G) A part of the Dimerizing Loop (332-342) is fitted in the cryo-EM density. (H) Two β-strands of the ABH domain (79-88 and 102-111) are fitted in the cryo-EM density. (I) A C-terminus of the ABH domain (509-515) is fitted in the cryo-EM density.
