## Supplementary material for "Structure determinants of DANGEROUS MIX 3, an alpha/beta hydrolase, for triggering NLR-mediated genetic incompatibility in plants": Figure S3

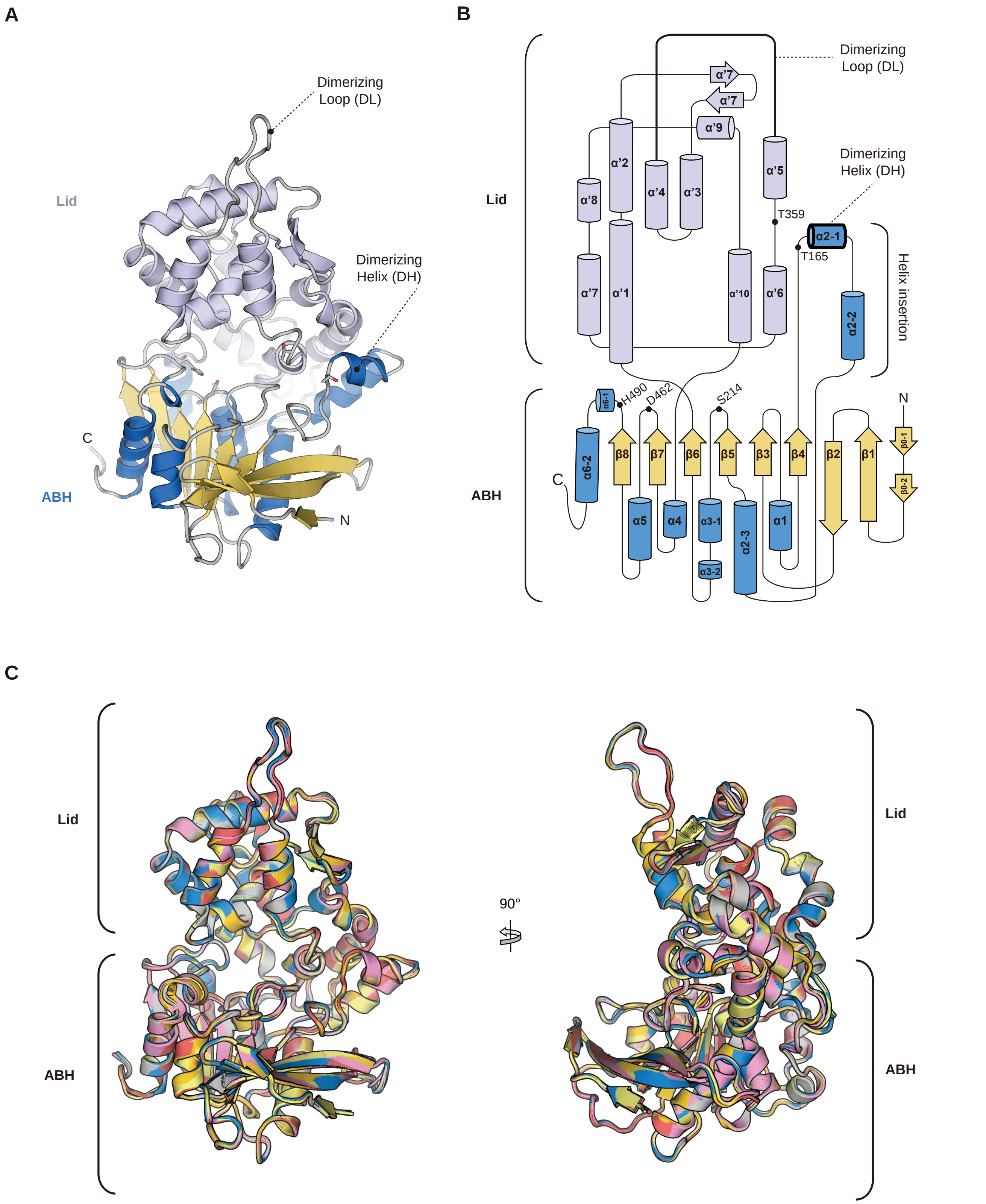


**Figure S3. Topology diagram of ${}_{\Delta TP}DM3^{Col\text{-}0}$ monomer and comparison of individual monomer subunits of ${}_{\Delta TP}DM3^{Col\text{-}0}$ hexamer. Related to Figure 2.** (A) The cryo-EM structure of ${}_{\Delta TP}DM3^{Col\text{-}0}$ monomer is shown. The secondary structure of the lid domain is colored in pale purple. α-helix and β-strand of the ABH domain are colored blue and yellow respectively. (B) The topology diagram of ${}_{\Delta TP}DM3^{Col\text{-}0}$ monomer shows arrangement of the secondary structures. The secondary structure of the lid domain is labeled with apostrophe ('). Positions of catalytic triad (S214, D462 and H490) amino acids and the polymorphic residues (T165 and T359) are denoted as black dots. DH, DL and helix insertion are labeled accordingly. (C) Six ${}_{\Delta TP}DM3^{Col\text{-}0}$ monomers are superimposed and shown as a cartoon. RMSD values between ${}_{\Delta TP}DM3^{Col\text{-}0}$ subunits are less than 0.286 Å.
