## Supplementary material for "Structure determinants of DANGEROUS MIX 3, an alpha/beta hydrolase, for triggering NLR-mediated genetic incompatibility in plants": Figure S4

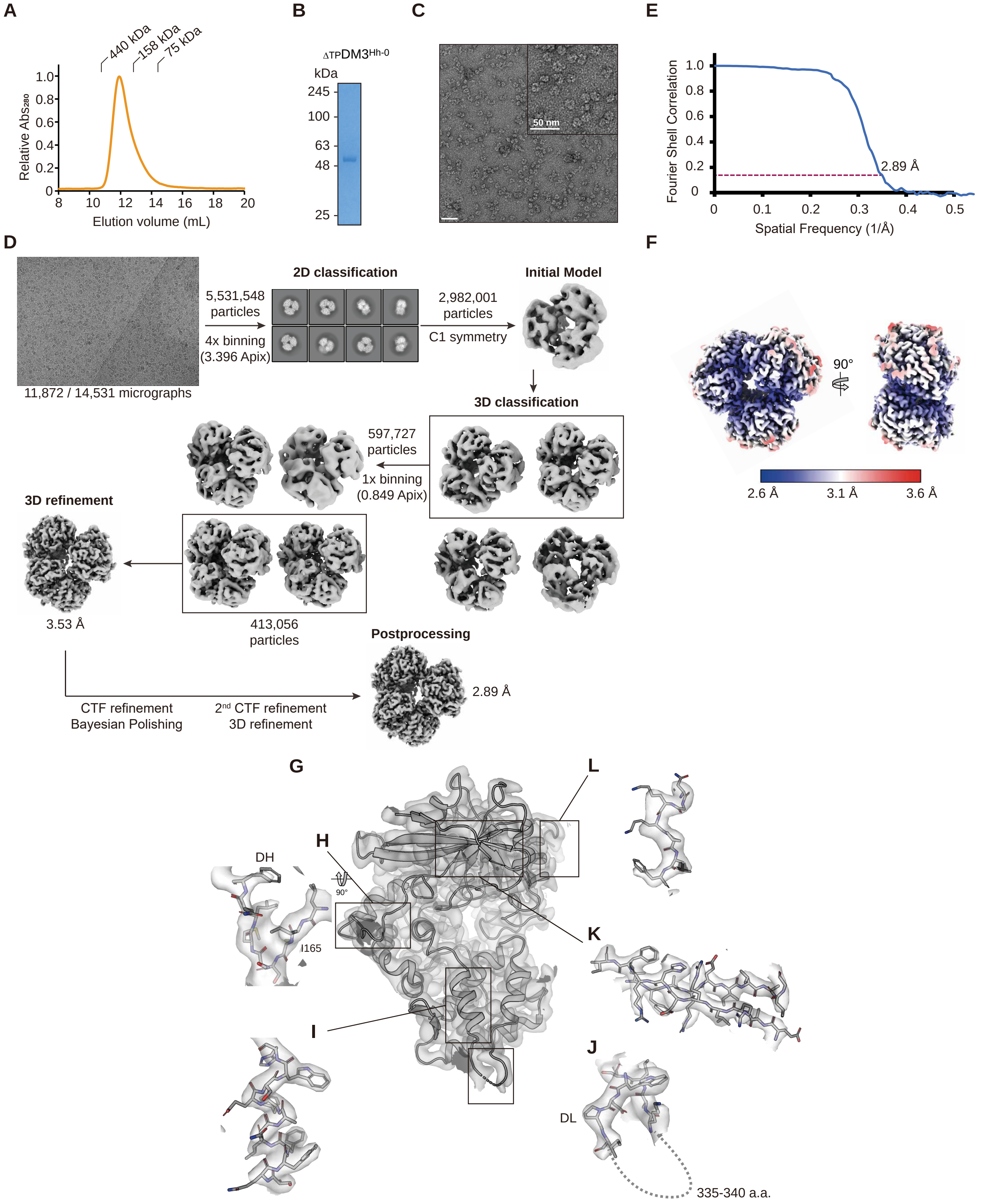


**Figure S4. Biochemical characterization, cryo-EM image processing and validation of ΔTPDM3$^{Hh-0}$. Related to Figure 4.** (A) A gel-filtration chromatogram of the purified recombinant ΔTPDM3$^{Hh-0}$ is shown. (B) 1 μg of the purified ΔTPDM3$^{Hh-0}$ protein was examined by SDS-PAGE. (C) A negative-stain EM shows ΔTPDM3$^{Hh-0}$ oligomer. An enlarged micrograph is shown with the scale bar (50 nm). (D) A flowchart of the ΔTPDM3$^{Hh-0}$ cryo-EM data processing shows a representative micrograph, 2D class averages and a final EM map (bottom right). Detailed descriptions of the data processing is in STAR Methods. (E) FSC curve shows resolution of the EM map of ΔTPDM3$^{Hh-0}$ as 2.89 Å with 0.143 criteria. (F) Local resolution of the cryo-EM map of DM3$^{Hh-0}$ is estimated and colored by Relion. The scale bar is depicted at the bottom. (G) Cryo-EM structure of ΔTPDM3Hh-0 monomer is fitted in the cryo-EM map. The monomer structure and the cryo-EM map are colored dark grey and grey respectively. (H) The melted DH of the dimer interface of DM3$^{Hh-0}$ (164-172) is fitted in the cryo-EM density. I165 is located before the DH (DH; 166-171). (I) An alpha helix of lid domain (344-355) is fitted in the cryo-EM density. (J) DL (331-344 ; 335-340 are disordered) is fitted in the cryo-EM density. The cryo-EM density for residues 335 and 340 was insufficient to model the structure of DM3Hh-0. (K) Two β-strands of the ABH domain (79-88 and 102-111) are fitted in the cryo-EM density. (L) A C-terminus of the ABH domain (509-515) is fitted in the cryo-EM density.
