## Supplementary material for "Structure determinants of DANGEROUS MIX 3, an alpha/beta hydrolase, for triggering NLR-mediated genetic incompatibility in plants": Figure S5

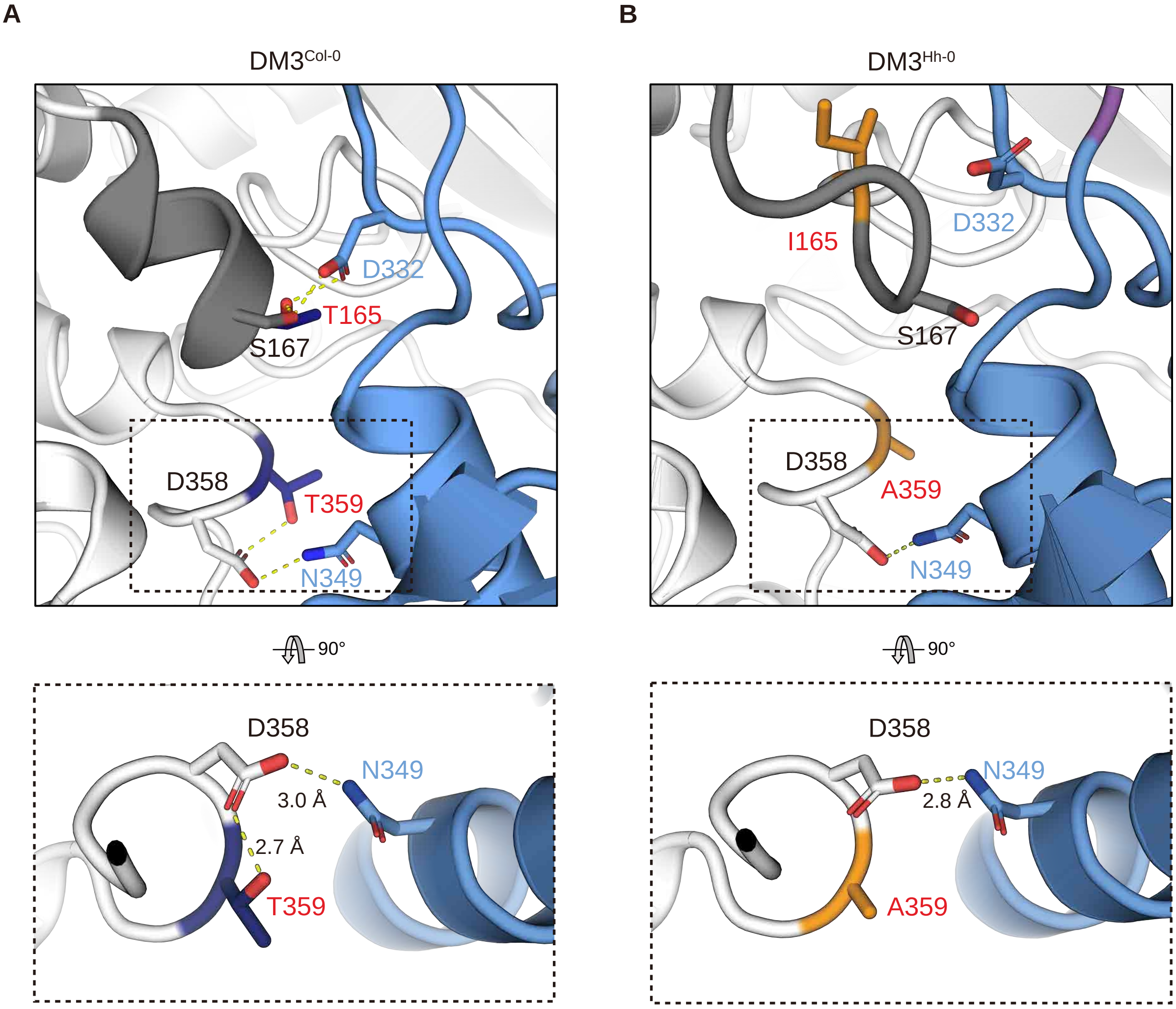


**Figure S5. The T359A polymorphism of DM3$^{Hh\text{-}0}$ does not affect the structure of the dimer interface. Related to Figure 4.** (A) The dimer interface of DM3$^{Col\text{-}0}$ shows T165 and T359 (navy) and its interacting amino acids. Electrostatic interactions between amino acids are represented as dotted lines. Dimer interface containing T359 is outlined in a box and shown at the bottom. (B) The dimer interface of DM3$^{Hh\text{-}0}$ shows I165 and A359 (orange) and with its interacting amino acids. Electrostatic interactions between amino acids are represented as dotted lines. Dimer interface containing A359 is outlined in a box and shown at the bottom.
