## Supplementary material for "Structure determinants of DANGEROUS MIX 3, an alpha/beta hydrolase, for triggering NLR-mediated genetic incompatibility in plants": Figure S6

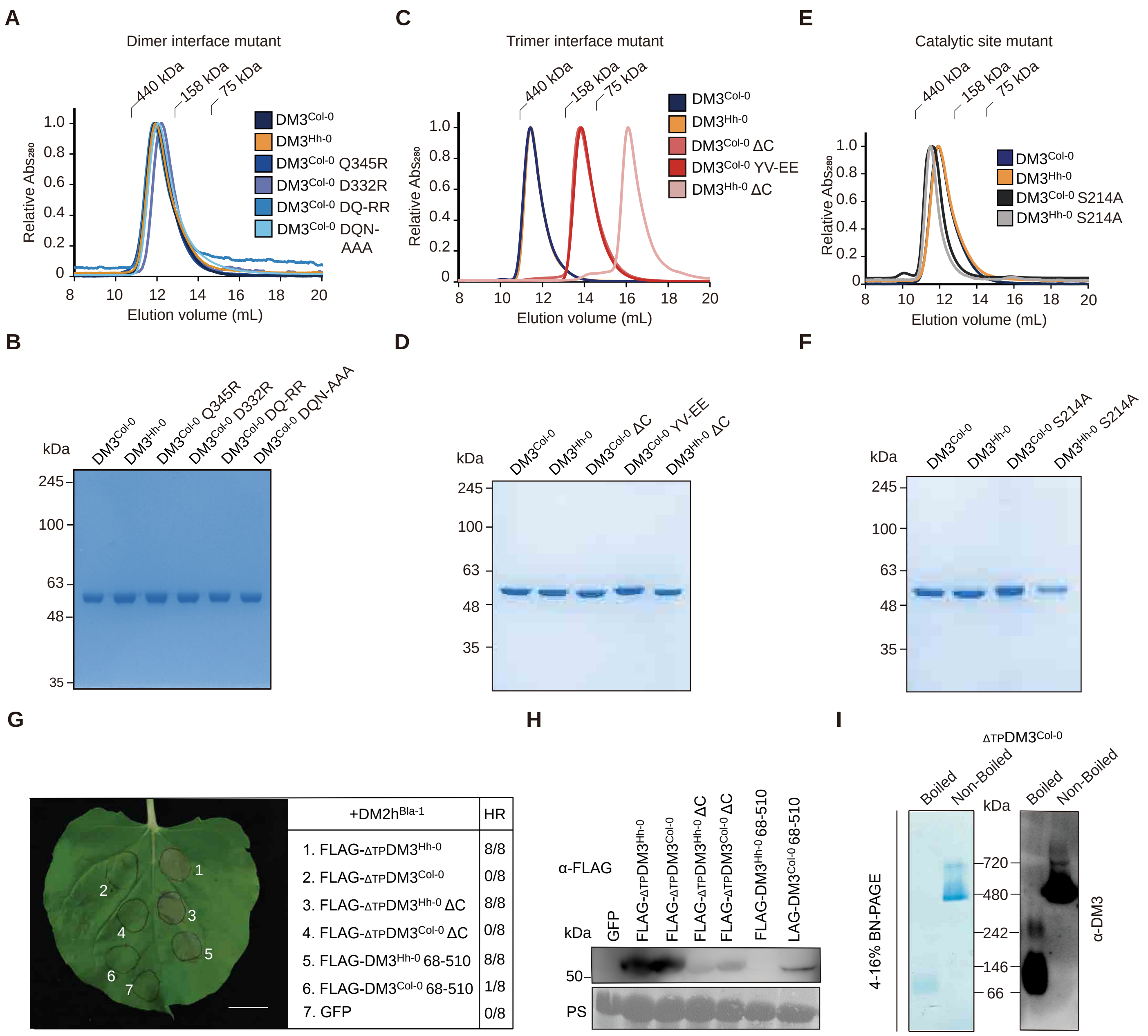


**Figure S6. Purification of recombinant structure-guided mutant DM3 proteins for biochemical analysis and the smallest unit of DM3. Related to Figure 4, 5, 6.** (A) A gel-filtration chromatogram of dimer interface mutants (DQ-RR; D332R Q345R and DQN-AAA; D332A Q345A N349A) and wild type $_{\Delta TP}$DM3$^{Col-0}$ and $_{\Delta TP}$DM3$^{Hh-0}$ are shown. The standard molecular weight marker is labeled on top. (B) An SDS-PAGE verifies purification of recombinant dimer interface mutation of $_{\Delta TP}$DM3$^{Col-0}$. 1 µg of purified protein was loaded on each well. (C) A gel-filtration chromatogram of trimer interface mutants (ΔC; Δ511-515 and YV-EE; Y460E V484E) and wild type $_{\Delta TP}$DM3$^{Col-0}$ and $_{\Delta TP}$DM3$^{Hh-0}$ are shown. (D) An SDS-PAGE verifies the purity of recombinant trimer interface mutants. 1 µg of purified protein was loaded on each well. (E) A gel-filtration chromatogram of $_{\Delta TP}$DM3$^{Col-0}$ and $_{\Delta TP}$DM3$^{Hh-0}$ catalytic site mutants (S214A) and wild type $_{\Delta TP}$DM3$^{Col-0}$ and $_{\Delta TP}$DM3$^{Hh-0}$ are shown. (F) An SDS-PAGE verifies the purity of recombinant catalytic site mutants. (G) Expression of FLAG-tagged truncated DM3 with DM2h$^{Bla-1}$ in *Nb*. DM2h$^{Bla-1}$ was driven by endogenous promoter, while DM3s and GFP were driven by 35S promoter. Photos were taken five days after infiltration. The numbers indicate leaves with fully developed HR out of eight leaves. Scale bar equals 2 cm. (H) Western blot of truncated DM3. Expression of all genes were was driven by 35S promoter. Agroinfiltrated *Nb* leaf samples were collected at 30 hours after infiltration. PS: Ponceau S. (I) BN-PAGE of $_{\Delta TP}$DM3$^{Col-0}$ recombinant protein. DM3 protein was loaded with or without 95°C 5 minutes treatment, 1 µg for coomassie blue staining and 1 ng for western blot.
