## Supplementary material for "Structure determinants of DANGEROUS MIX 3, an alpha/beta hydrolase, for triggering NLR-mediated genetic incompatibility in plants": Figure S7

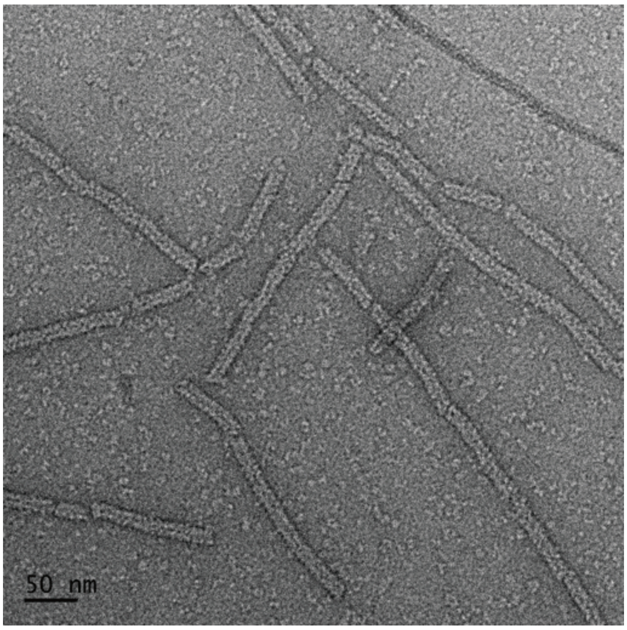


**Figure S7. Negative stain electron microscopy image of the trimer interface mutant $_{\Delta TP}DM3^{Col\text{-}0}$ ΔC.** Imaging was performed using 0.03 mg/mL of protein. The micrograph shows filamentous DM3 structures, with a 50 nm scale bar.
