## Supplementary material for "Structure determinants of DANGEROUS MIX 3, an alpha/beta hydrolase, for triggering NLR-mediated genetic incompatibility in plants": Table S1, Table S2

**Supplemental Information**

| | $_{\Delta TP}DM3^{Col\text{-}0}$ | $_{\Delta TP}DM3^{Hh\text{-}0}$ |
|---|---|---|
| **Sample Preparation** | | |
| Grid | Quantifoil R1.2/1.3 300 | Quantifoil R1.2/1.3 300 |
| Treatment on grid | Graphene oxide | Graphene oxide |
| Cryo-specimen freezing | Vitrobot Mark IV | Vitrobot Mark IV |
| **Data collection and processing** | | |
| Electron Microscope | Glacios (200 keV) | TFS Krios G4 |
| Detecting device | Falcon III | Gatan K3 BioQuantum |
| Acquisition mode | Electron counting | Electron counting (CDS) |
| Pixel size (Å/pix) | 0.8943 | 0.849 |
| Electron exposure ($e^-/Å^2$) | 40 | 49.8 |
| Number of frames (no.) | 40 | 48 |
| Defocus range (µm) | -0.8 ~ -2.0 | -0.9 ~ -2.3 |
| Symmetry imposed | C1 | C1 |
| Initial / Final particle used (no.) | 1,845,172 / 330,509 | 5,531,548 / 413,056 |
| Resolution (Å) | 3.39 | 2.89 |
| FSC threshold | 0.143 | 0.143 |
| Applied B-factor ($Å^{\wedge 2}$) | -163.287 | -20 |
| **Model composition** | | |
| Nonhydrogen atoms | 21,360 | 21,120 |
| Protein residues | 2,688 | 2,652 |
| **R.m.s. deviations** | | |
| Bond length (Å) | 0.012 | 0.012 |
| Bond angles (°) | 1.924 | 1.842 |
| **Validation** | | |
| MolProbity score | 1.04 | 0.75 |
| Rotamer outliers (%) | 0 | 0.18 |
| Clashscore (%) | 1.53 | 0.79 |
| C-beta outliers (%) | 0 | 0 |
| Mask CC | 0.72 | 0.82 |
| **Ramachandran Plot** | | |
| Favored (%) | 97.23 | 98.10 |
| Allowed (%) | 2.77 | 1.90 |
| Outliers (%) | 0 | 0 |

**Table S1. Cryo-EM sample preparation, data collection, processing, and model validation statistics (related to Figures 2 and 4, STAR Methods).**

| Primer | Purpose | Sequence 5'-3' |
|---|---|---|
| DM3 53F | $_{\Delta TP}$DM3 cloning into entry vector for plant expression | CAGGCTTTCGAATTCCAATGGAAGCGGGATCGGTTTACG |
| DM3 515R | $_{\Delta TP}$DM3 cloning into entry vector for plant expression | TGGGTCTAGATAGGAATTCGGTCAAAAGAGAGGCTTTTCCCATTG |
| DM3 D332R F | DM3 mutagenesis PCR in entry vector | GGAGAGAGTATGGCGTCCTATTTTAGTTACTGGAG |
| DM3 D332R R | DM3 mutagenesis PCR in entry vector | CTCCAGTAACTAAAATAGGACGCCATACTCTCTCC |
| DM3 Q345R F | DM3 mutagenesis PCR in entry vector | CCAAAGTGTATTAGTCGGTTCTTCTTAAACGCTG |
| DM3 Q345R R | DM3 mutagenesis PCR in entry vector | CAGCGTTTAAGAAGAACCGACTAATACACTTTGG |
| DM3 DQN-AAA F | DM3 mutagenesis PCR in entry vector | GTTACTGGAGCTCCAAAGTGTATTAGTGCGTTCTTCTTAGCCGCTGTAAG |
| DM3 DQN-AAA R | DM3 mutagenesis PCR in entry vector | CTAATACACTTTGGAGCTCCAGTAACTAAAATAGGAGCCCATACTC |
| DM3 V484E F | DM3 mutagenesis PCR in entry vector | CAGACTTTGGGAAACGAATGAGTTTATGCATTCGGG |
| DM3 V484E R | DM3 mutagenesis PCR in entry vector | CCCGAATGCATAAACTCATTCGTTTCCCAAAGTCTG |
| DM3 Y460E R | DM3 mutagenesis PCR in entry vector | CATACATATCTTCTTCGTAAACCGCTGCAG |
| DM3 Y460E F | DM3 mutagenesis PCR in entry vector | CTGCAGCGGTTTACGAAGAAGATATGTATG |
| DM3 510 R | $_{\Delta TP}$DM3 ΔC cloning into entry vector for plant expression | CCATGGATCCTCACCCATTGATCATTCCCAACAAATG |
| DM3Col0_52F_SalI_pET28aIN | $_{\Delta TP}$DM3 cloning into pET28a vector | ATATGTCGACAAGCTGAAGCGGGATCGGTTTAC |

| | | |
|---|---|---|
| DM3_515R_Hind III | $_{\Delta TP}$DM3 cloning into pET28a vector | ATATAAGCTTTCAAAAGAGAGGCTTTTTCCCATTG |
| DM3_Q345A_N349A_F | DM3 mutagenesis PCR in pET28a vector | CCAAAGTGTATTAGTGCGTTCTTCTTAGCGGCTTTTGAG |
| DM3_Q345A_N349A_R | DM3 mutagenesis PCR in pET28a vector | CTCAAAAGCCGCTAAGAAGAACGCACTAATACACTTTGG |
| DM3_D332A_F | DM3 mutagenesis PCR in pET28a vector | GTTGGAGAGAGTATGGGCGCCTATTTTAGTTAC |
| DM3_D332A_R | DM3 mutagenesis PCR in pET28a vector | GTAACTAAAATAGGCGCCCATACTCTCTCCAAC |
| DM3_D332R_F | DM3 mutagenesis PCR in pET28a vector | GGAGAGAGTATGGCGTCCTATTTTAGTTACTGG |
| DM3_D332R_R | DM3 mutagenesis PCR in pET28a vector | CCAGTAACTAAAATAGGACGCCATACTCTCTCC |
| DM3_Q345R_F | DM3 mutagenesis PCR in pET28a vector | GGAGCTCCAAAGTGTATTAGTCGTTTCTTCTTAAACGC |
| DM3_Q345R_R | DM3 mutagenesis PCR in pET28a vector | GCGTTTAAGAAGAAACGACTAATACACTTTGGAGCTCC |
| DM3_511TER_F | DM3 mutagenesis PCR in pET28a vector | GGGAATGATCAATGGGTGAAAGCCTCTCTTTTG |
| DM3_511TER_R | DM3 mutagenesis PCR in pET28a vector | CAAAAGAGAGGCTTTCACCCATTGATCATTCCC |
| DM3_Y460E_F | DM3 mutagenesis PCR in pET28a vector | GCTGCAGCGGTTTACGAAGAAGATATGTATG |
| DM3_Y460E_R | DM3 mutagenesis PCR in pET28a vector | CATACATATCTTCTTCGTAAACCGCTGCAGC |
| DM3_V484E_F | DM3 mutagenesis PCR in pET28a vector | CGGGTATCAGACTTTGGGAAACGAATGAGTTTATGC |
| DM3_V484E_R | DM3 mutagenesis PCR in pET28a vector | GCATAAACTCATTCGTTTCCCAAAGTCTGATACCCG |
| DM3_S214A_F | DM3 mutagenesis PCR in pET28a vector | GGACAATTTTGGGTCAGGCGTTCGGTGGCTTTTGTGC |

| | | |
|---|---|---|
| DM3_S214A_R | DM3 mutagenesis PCR in pET28a vector | GCACAAAAGCCACCGAACGCCTGACCCAAAATTGTCC |

**Table S2. Primer list (related to STAR Methods)**
